# BK channel properties correlate with neurobehavioral severity in three *KCNMA1*-linked channelopathy mouse models

**DOI:** 10.1101/2022.02.15.478992

**Authors:** Su Mi Park, Cooper E. Roache, Philip H. Iffland, Hans J. Moldenhauer, Katia K. Matychak, Amber E. Plante, Abby G. Lieberman, Peter B. Crino, Andrea L. Meredith

**Affiliations:** Departments of Physiology, University of Maryland School of Medicine, Baltimore, MD, 21201; Departments of Pharmacology, University of Maryland School of Medicine, Baltimore, MD, 21201; Departments of Neurology, University of Maryland School of Medicine, Baltimore, MD, 21201

**Author notes:** Corresponding author: Andrea L. Meredith, Ph.D, Dept. of Physiology, University of Maryland School of Medicine 655 W. Baltimore St., Baltimore, MD 21201) 706-5991.

**Keywords:** BK channel, KCa1.1, calcium-activated potassium channel, *Kcnma1*, *Slo*, *slowpoke*, seizure, epilepsy, hippocampus, dentate gyrus, dyskinesia, paroxysmal nonkinesigenic dyskinesia, PNKD, PNKD3, movement disorder

## Abstract

*KCNMA1* forms the pore of BK K^+^ channels, which regulate neuronal and muscle excitability. Recently, genetic screening identified heterozygous *KCNMA1* variants in a subset of patients with debilitating paroxysmal non-kinesigenic dyskinesia, presenting with or without epilepsy (PNKD3). However, the relevance of *KCNMA1* mutations and the basis for clinical heterogeneity in PNKD3 has not been established. Here we evaluate the relative severity of three *KCNMA1* patient variants in BK channels, neurons, and mice. In heterologous cells, BK^N999S^ and BK^D434G^ channels displayed gain-of-function (GOF) properties, whereas BK^H444Q^ channels showed loss-of-function (LOF) properties. The relative degree of channel activity was BK^N999S^ > BK^D434G^ > WT > BK^H444Q^. BK currents and action potential firing were increased, and seizure thresholds decreased, in *Kcnma1*^N999S/WT^ and *Kcnma1*^D434G/WT^ transgenic mice but not *Kcnma1*^H444Q/WT^ mice. In a novel behavioral test for paroxysmal dyskinesia, the more severely affected *Kcnma1*^N999S/WT^ mice became immobile after stress, consistent with stress-induced immobility episodes observed in PNKD3-affected individuals. Homozygous *Kcnma1*^D434G/D434G^ mice showed similar immobility, but in contrast, homozygous *Kcnma1*^H444Q/H444Q^ mice displayed hyperkinetic behavior. These data establish the relative pathogenic potential of patient alleles as N999S > D434G > H444Q and validate *Kcnma1*^N999S/WT^ mice as a model for PNKD3 with increased seizure propensity.

## INTRODUCTION

*KCNMA1*-linked channelopathy encompasses an array of neurological symptoms associated with clinical detection of a *KCNMA1* variant. Affected individuals typically present with epilepsy and/or dyskinesia, but also have other types of movement disorder including ataxia, developmental delay, intellectual disability, and brain and structural abnormalities (Bailey *et al*., 2019; Liang *et al*., 2019; Miller *et al*., 2021). The basis for these symptoms is not mechanistically established but is likely similar to other neurological channelopathies involving direct or indirect changes in neuronal excitability leading to excitation-inhibition imbalance (Benatar, 2000; Menezes *et al*., 2020). *KCNMA1* genotype-phenotype correlation is an active area of investigation with >40 variants identified in this patient population to date (Miller *et al*., 2021 and ALM unpublished data). Since most variants arise *de novo* in a single heterozygous proband, whether ‘*KCNMA1* channelopathy’ is a *bona fide* monogenic disorder, or results from intergenic and developmental interactions, is not well-understood. Animal models for the most common variants are needed to validate genotype-phenotype associations and to investigate disease mechanisms and manifestations over lifespan.

*KCNMA1* encodes the ‘Big K^+’^ (BK) channel, activated by voltage and intracellular Ca^2+^ (Figure 1). BK currents are prominent in the central nervous system and smooth muscle (Bailey *et al*., 2019; Contet *et al*., 2016; Latorre *et al*., 2017). Neuronal BK channels regulate action potential repolarization and fast afterhyperpolarizations (fAHP) to set firing rates (Gu *et al*., 2007; Montgomery and Meredith, 2012; Sah and Faber, 2002; Shao *et al*., 1999) and neurotransmission (Golding *et al*., 1999; Raffaelli *et al*., 2004; Sailer *et al*., 2006). *KCNMA1* knockout mice (*Kcnma1*^—/—^) show prominent smooth muscle, neurobehavioral, and locomotor deficits, associated with widespread alterations in cellular excitability (MGI:99923; Bailey *et al*., 2019; Meredith *et al*., 2004; Sausbier *et al*., 2005a; Sausbier *et al*., 2004). However, *Kcnma1*^—/—^ mice do not phenotypically exhibit *KCNMA1*-linked channelopathy symptoms. Moreover, the largest cohort of clinically distinguishable patients harbor GOF, rather than LOF, alleles with respect to BK channel activity (Miller *et al*., 2021).

**Figure 1.**
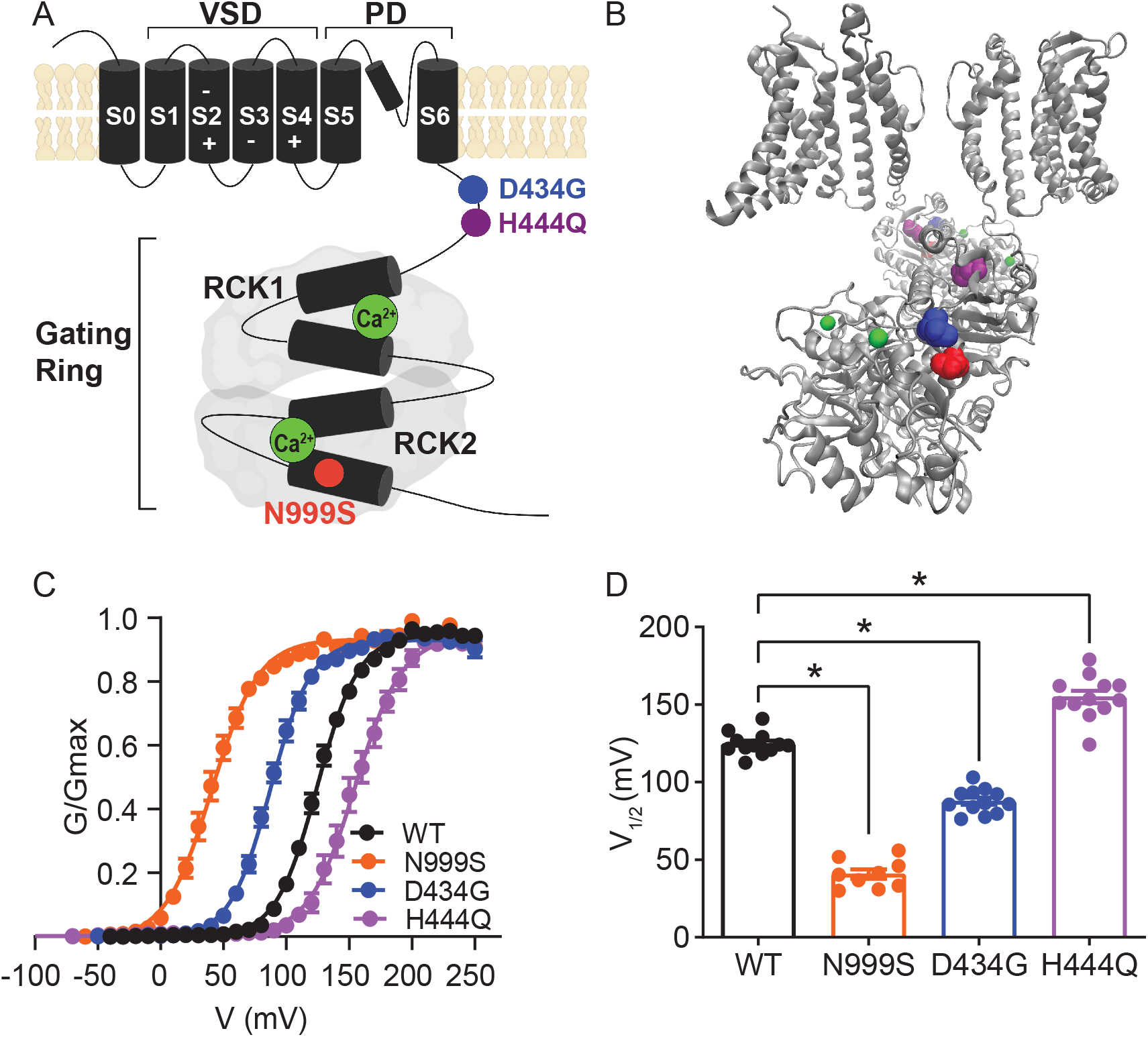
Location and consequence of *KCNMA1* variants in the BK K^+^ channel. (A) *KCNMA1* forms the homotetrameric BK channel. Each α subunit is comprised of seven transmembrane domains (S0‒S6) and an intracellular gating ring. Pore opening and closing is regulated by voltage sensitive residues in S2‒S4 and two Regulators of Conductance of Potassium (RCK) domains in the gating ring, each containing a Ca^2+^ binding site (green) (Yang *et al*. 2015, Giraldez and Rothberg 2017). H444Q (purple) and D434G (blue) are located within the βB‒αB and αA and βB of the AC domain, respectively, a region within RCK1 affecting Ca^2+^‒dependent gating (Du *et al*. 2005, Tao and MacKinnon 2019). N999S (red) is located at the helix bend in the middle of the S10 domain within RCK2 (Tao and MacKinnon 2019). (B) BK channel structure showing two opposing subunits including the voltage sensor domain (VSD), pore domain (PD), and gating ring with Ca^2+^ bound (PDB 6V38). (C) Inside‒out patch‒clamp recordings from BK^WT^, BK^N999S^, BK^D434G^, BK^H444Q^ channels expressed in HEK293 cells. Macroscopic BK currents were activated with the voltage protocol described in Supplemental Figure S1 in symmetrical K^+^ and 1 μM intracellular Ca^2+^. Normalized conductance‒voltage (G‒V) relationships fit with Boltzmann functions (solid lines). There was no change in the slope factor (z) for any of the variants (*p* = 0.06, One-way ANOVA). BK^WT^ (n=12), BK^N999S^ (n=9), BK^D434G^ (n=12), and BK^H444Q^ (n=12). (D) Voltage of half‒maximal activation (V1/2) obtained from Boltzmann fits for individual patches. \**p* < 0.0001. One-way ANOVA with Dunnett’s post hoc.

Two GOF *KCNMA1* variants, D434G and N999S, account for half of the patient population (Bailey *et al*., 2019; Miller *et al*., 2021). Both variants cause BK channel activation at more negative membrane potentials, speed activation, and slow deactivation (Diez-Sampedro *et al*., 2006; Du *et al*., 2005; Li *et al*., 2018; Moldenhauer *et al*., 2020; Wang *et al*., 2009; Yang *et al*., 2010). The majority of individuals harboring D434G and N999S variants present with paroxysmal non-kinesigenic dyskinesia (PNKD type 3; OMIM #609446), characterized by varying degrees of dystonia, hypotonia, or non-narcoleptic cataplexy. PNKD3 episodes occur with short duration and high frequency, often hundreds of times per day (Du *et al*., 2005; Heim *et al*., 2020; Keros *et al*., 2022; Li *et al*., 2018; Wang *et al*., 2017; Zhang *et al*., 2015). Just under half of patients experience seizure of varying types, including absence, atonic, myoclonic, and generalized tonic-clonic (GTC). However, epilepsy and PNKD are not consistently co-morbid (Du *et al*., 2005; Miller *et al*., 2021). Individuals with putative LOF variants report additional movement disorders including dyskinesia, axial hypotonia, tremor, or ataxia, in addition to various seizure types (Du *et al*., 2020; Liang *et al*., 2019; Rodrigues Bento *et al*., 2021; Tabarki *et al*., 2016; Yesil *et al*., 2018). It is not yet clear whether variations in symptomatic presentation result from incomplete or inconsistent clinical evaluations, or genuine genotype-phenotype differences within either GOF or LOF cohorts.

We address these questions through heterologous, neuronal, and neurobehavioral validation for three patient-associated *KCNMA1* variants in mouse models. The GOF BK^N999S^ and BK^D434G^ channels produced increased neuronal BK currents and firing as heterozygous alleles in transgenic mice, while heterozygous LOF BK^H444Q^ channels were insufficient to alter neuronal properties. Mice were evaluated in a series of spontaneous and evoked seizure and locomotor assays. N999S propagated the largest symptomatic burden with chemoconvulsant challenge and stress-triggered dyskinesia, supporting the conclusion that this variant has the greatest monogenic pathogenicity, followed by D434G, *Kcnma1*^—/—^, and H444Q. The results identify *Kcnma1*^N999S/WT^ mice as a PNKD3 model with the highest phenotypic similarity to patients harboring *KCNMA1* GOF variants. Our findings further establish the stress-induced PNKD assay to delineate distinct symptomatic manifestations between GOF and LOF alleles, supporting its utility in a battery of neurobehavioral evaluations to define *KCNMA1*-linked channelopathy models.

## RESULTS

### Patient variants confer GOF (N999S and D434G) and LOF (H444Q) properties on BK channel activity

A comparative assessment for three dyskinesia-associated patient variants (N999S, D434G, and H444Q) was performed within the human BK channel (Figure 1). BK channel function was assessed using inside-out patch-clamp recordings in HEK293 cells (Figure 1C-D, S1A). Patches from cells expressing BK^WT^, BK^N999S^, BK^D434G^, BK^H444Q^ channels were activated with depolarizing voltage steps, and the voltage-dependence of activation and kinetics were assessed from macroscopic currents (Figure 1 and S1). Conductance versus voltage (G-V) relationships (Figure 1C) were assessed by the voltage of half-maximal activation (V1/2; Figure 1D).

BK^WT^ currents had a V1/2 of 125 ± 2 mV. Introduction of N999S and D434G mutations shifted the G-V relationships to more negative membrane potentials (V1/2: BK^N999S^ 41 ± 3 mV and BK^D434G^ 88 ± 2 mV), confirming their GOF effect at all voltages. The decrease in V1/2 for BK^N999S^ channels compared to BK^WT^ was 20-30 mV larger than observed in prior studies with different splice variants and intracellular Ca^2+^ (Li *et al*., 2018; Moldenhauer *et al*., 2020). Here under equivalent conditions, N999S produced a larger hyperpolarizing shift from WT (ΔV1/2 = 84 mV) versus D434G (ΔV1/2 = 37 mV). In addition, N999S and D434G produced faster BK channel activation and slower deactivation compared to WT (Figure S1). Altogether, BK^N999S^ channels showed greater GOF properties than BK^D434G^ in all parameters under these conditions, corroborating the relative severity predicted from prior work.

In contrast, introduction of the H444Q variant shifted the G-V relationship to more positive potentials (V1/2: BK^H444Q^ 155 ± 4 mV), consistent with LOF effects. H444Q produced changes in channel opening and closing further consistent with LOF effects, slowing activation and speeding deactivation (Figure S1). H444Q produced a smaller difference from WT than either GOF variant (ΔV1/2 = 30 mV), identifying H444Q as a comparatively mild variant. The results indicate that N999S produces the strongest effect on BK channel activation in the GOF direction, followed by D434G (GOF), and H444Q (LOF).

### Generation of N999S, D434G, and H444Q mouse models

Correlation between patient genotype and phenotype has only been established for a single *KCNMA1* variant so far, D434G, an autosomal dominant that co-segregates with PNKD and epilepsy in a multi-generation pedigree (Du *et al*., 2005). D434G pathogenicity is further corroborated by mouse and fly models, which show alterations in neuronal excitability, brain and motor function (Dong *et al*., 2021; Kratschmer *et al*., 2021). In contrast, N999S and H444Q lack this direct evidence due to the absence of familial transmission among the children that carry these variants (Miller *et al*., 2021). N999S is the most common *de novo KCNMA1* variant (∼17% of all patients), found as heterozygous in every case. About half of individuals harboring N999S alleles are diagnosed with seizure, PNKD, or both (Keros *et al*., 2022; Li *et al*., 2018; Wang *et al*., 2017; Zhang *et al*., 2015), suggesting a strong potential to be causative in channelopathy symptoms. H444Q is found in a single case and is one of several putative LOF variants where affected individuals have dyskinesia-like paroxysms (Miller *et al*., 2021). This proband had a history of abnormal EEG, unresolved with respect to the diagnosis of epilepsy, but also harbors three additional genetic findings (ALM unpublished data).

To establish genotype-phenotype correlations, heterozygous mice replicating the patient genotypes were first evaluated. Each variant was introduced as a single nucleotide mutation into the mouse *Kcnma1* gene using CRISPR base-editing (Figure S2). In all animal experiments, investigators were blinded to genotype during data collection and analysis, and WT controls were compared to transgenic littermates within individual transgenic lines. *Kcnma1*^N999S/WT^, *Kcnma1*^D434G/WT^, and *Kcnma1*^H444Q/WT^ mice were grossly behaviorally and morphologically normal with no notable spontaneous paroxysms, gait abnormalities, or visually detectable seizures during home cage observation. *Kcnma1*^D434G/WT^ and *Kcnma1*^H444Q/WT^ intercrosses produced homozygous progeny that were also visually normal. However, *Kcnma1*^N999S/WT^ intercrosses produced no homozygous pups (see Methods). The absence of homozygous N999S progeny is similar to *Tg-BK^R207Q^* mice harboring another strong GOF mutation that showed lethality in the homozygous allele configuration (Montgomery and Meredith, 2012). Given the inability to generate homozygous N999S mice, gene expression was analyzed from hippocampus and cerebellum of *Kcnma1*^N999S/WT^ and WT littermates (n = 3 mice each genotype and tissue). No significant differences were found in the levels of *Kcnma1* (1.07-fold change, P = 0.79, FDR = 0.99, ANOVA with eBayes test), or BKβ subunits expressed in brain: *Kcnmb1* (−1.07, P = 0.14), *Kcnmb2* (1.00, P = 0.84), and *Kcnmb4* (−1.03, P = 0.46).

### N999S and D434G increase BK current in hippocampal neurons

From heterologous cells, we predicted that the variants would have a strong (N999S), intermediate (D434G), or weak (H444Q) potential to alter neuronal BK current levels in transgenic mice. However, heterozygous patient genotypes create the possibility for hetero- tetramer channel formation (Geng *et al*., 2021), necessitating understanding the relative GOF and LOF effects *in vivo* from BK current levels in heterozygous transgenics compared to WT littermates. Recordings were made in the dentate gyrus of the hippocampus, where BK channels are highly expressed, regulate neuronal excitability, and where changes in BK channel properties are associated with seizure (Kaufmann *et al*., 2010; Knaus *et al*., 1996; Misonou *et al*., 2006; Sailer *et al*., 2006; Sausbier *et al*., 2005b; Trimmer, 2015). In dentate granule cells, excitability is sensitive to changes in BK current in both directions, assessed using pharmacological inhibition as well as deletion of the β4 subunit (Brenner *et al*., 2005; Mehranfard *et al*., 2014; 2015). Loss of β4 creates GOF BK channels by speeding BK channel kinetics, and β4^—/—^ knockout mice have temporal lobe seizures (Jaffe and Brenner, 2018; Petrik *et al*., 2011; Wang *et al*., 2016; Whitmire *et al*., 2017).

BK currents from *Kcnma1*^WT/WT^ neurons activated at ‒40 mV, increasing to 21‒28 pA/pF at the highest voltage across mouse strains (Figure 2A-C). *Kcnma1*^N999S/WT^ neurons had a 69% increase in BK current compared to WT littermates (*Kcnma1*^WT/WT^ 13.0 ± 2.0 pA/pF and *Kcnma1*^N999S/WT^ 22.0 ± 1.8 pA/pF at ‒10 mV). The increased BK current likely results from alterations in BK channel activity, since *KCNMA1* expression was not changed in *Kcnma1*^N999S/WT^ neurons. *Kcnma1*^D434G/WT^ BK currents were 73% larger (*Kcnma1*^WT/WT^ 12.1 ± 2.6 pA/pF and *Kcnma1*^D434G/WT^ 20.9 ± 3.2 pA/pF at ‒10 mV), although not statistically different at any voltage due to variability. However, two copies of the D434G variant (*Kcnma1*^D434G/D434G^) resulted in the largest increase in BK current across voltages from ‒40 mV to the maximum (203%; 36.7 ± 5.9 pA/pF ‒10 mV). Interestingly, by direct comparison *Kcnma1*^D434G/WT^ BK current levels were similar to *Kcnma1*^N999S/WT^, despite the more severe phenotype for BK^N999S^ channels in heterologous cells (Figure 1).

**Figure 2.**
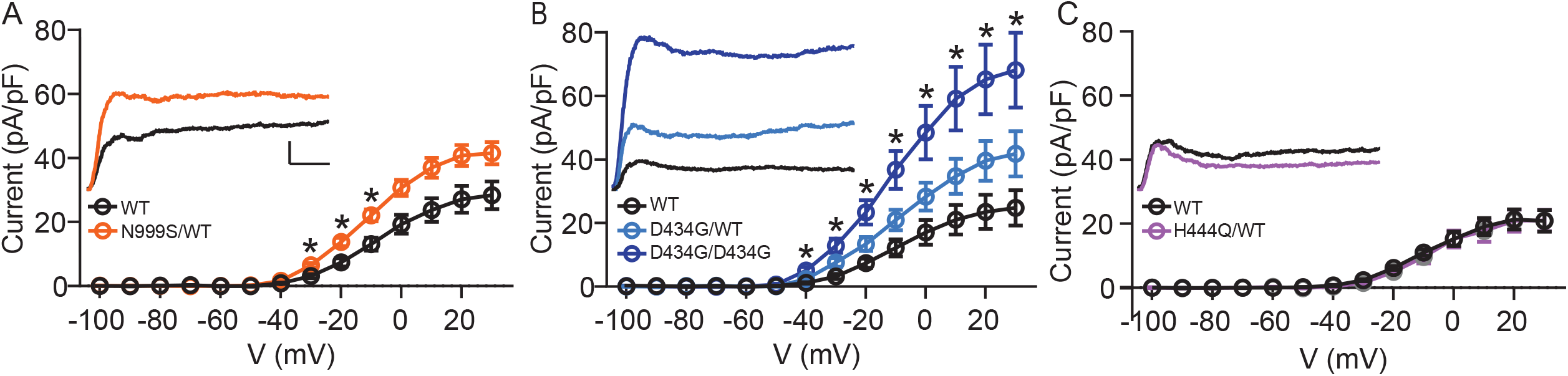
Increased BK current in *Kcnma1*^N999S/WT^ and *Kcnma1*^D434G/D434G^ granule neurons. Whole‒cell macroscopic BK currents were recorded in 1μM tetrodotoxin (TTX) and 2 mM 4- aminopyridine (4-AP), isolated with 10 μM paxilline, and normalized to cell capacitance. Activating voltage steps were applied from Vh of ‒90 mV, stepping from ‒100 to +30 mV for 150 ms, and back to ‒90 mV for 130 ms. (A‒C) Peak BK current density versus voltage relationships. Data are presented as mean ± SEM. *, *p* < 0.05, Two-way repeated measures ANOVA with Bonferroni post-hoc. *Insets:* Representative BK current traces at 30mV. Scale bars: 500 pA, 5 ms. (A) BK current density was larger in *Kcnma1*^N999S/WT^ neurons (n =16 neurons, 5 mice) compared to *Kcnma1*^WT/WT^ (n =14 neurons, 4 mice) at ‒30 mV (*p*=0.0114), ‒20 (*p*=0.0210), ‒10 (*p*=0.0426) voltage steps (indicated with *). (B) BK current density was larger in *Kcnma1* ^D434G/D434G^ neurons (n =12 neurons, 3 mice) compared to *Kcnma1*^WT/WT^ (n =10 neurons, 4 mice) at density at ‒40 mV (*p*=0.0112), ‒30 (*p*=0.0026), ‒20 (*p*=0.0031), ‒10 (*p*=0.0038), 0 (*p*=0.0078), 10 (*p*=0.0068), 20 (*p*=0.0071), 30 (*p*=0.0088) voltage steps (*). *Kcnma1*^D434G/WT^ mice (n=9 neurons, 3 mice) had higher BK current density compared to *Kcnma1*^WT/WT^ at ‒30mV only (*p*=0.0321). (C) BK current density was not different in *Kcnma1*^H444Q/WT^ neurons (n =7 neurons, 2 mice) compared to *Kcnma1*^WT/WT^ (n =6 neurons, 3 mice).

In contrast, BK currents in *Kcnma1*^H444Q/WT^ neurons were not significantly different compared to WT littermates at any voltage (*Kcnma1*^WT/WT^ 10.9 ± 1.0 pA/pF and *Kcnma1*^H444Q/WT^ 9.5 ± 1.9 pA/pF at ‒10 mV; ‒13% change). This establishes an allelic series of *Kcnma1*^D434G/D434G^ >> *Kcnma1*^D434G/WT^ ≈ *Kcnma1*^N999S/WT^ > *Kcnma1*^H444Q/WT^ with respect to BK current magnitude and support the potential for N999S and D434G to cause neurobehavioral changes, potentially in a dose-dependent manner. The pathogenic potential for *Kcnma1*^H444Q/WT^ is less clear and may require additional factors or mechanisms to support pathogenicity (i.e., other Ca^2+^ conditions, cell types, or gene interactions).

### N999S and D434G increase intrinsic neuronal excitability

Intrinsic excitability was next assessed in dentate granule neurons as an independent validation for neuronal pathogenicity. Both GOF and LOF BK channel mutations have the ability to alter neuronal activity in either direction, depending on the context (Bailey *et al*., 2019; Brenner *et al*., 2005; Montgomery and Meredith, 2012; Gu *et al.,* 2007; Sausbier *et al*., 2004).

Dentate granule cell input-output firing relationships were assessed in current-clamp mode (Figure 3). Firing rates increased with current injection in each *Kcnma1*^WT/WT^ littermate control dataset, reaching a peak of ∼40 Hz between 240 and 260 pA and then decreasing with higher current injections (Figure 3A-B). *Kcnma1*^N999S/WT^ firing was greater than *Kcnma1*^WT/WT^ littermate neurons in several key places. First, across the whole current injection range, firing was significantly increased in the middle portion (160 to 240 pA), ranging from 25-30% higher than WT (Figure 3A*i*, B*i*). After reaching the maximum, the firing still decreased instead of remaining higher through the full range of current injections. In addition, the initial slope of firing (0 to 160 pA) was greater in *Kcnma1*^N999S/WT^ neurons (0.22 ± 0.01 Hz/pA) compared to *Kcnma1*^WT/WT^ (0.18 ± 0.01 Hz/pA, Figure 3C*i*). Lastly, the maximal firing was 9.6 ± 1.8 Hz (125%) higher in *Kcnma1*^N999S/WT^ neurons versus *Kcnma1*^WT/WT^ (Figure 3D*i*). Taken together, *Kcnma1*^N999S/WT^ neurons respond to stimulation with higher firing and a shift in the input-output relationship.

**Figure 3.**
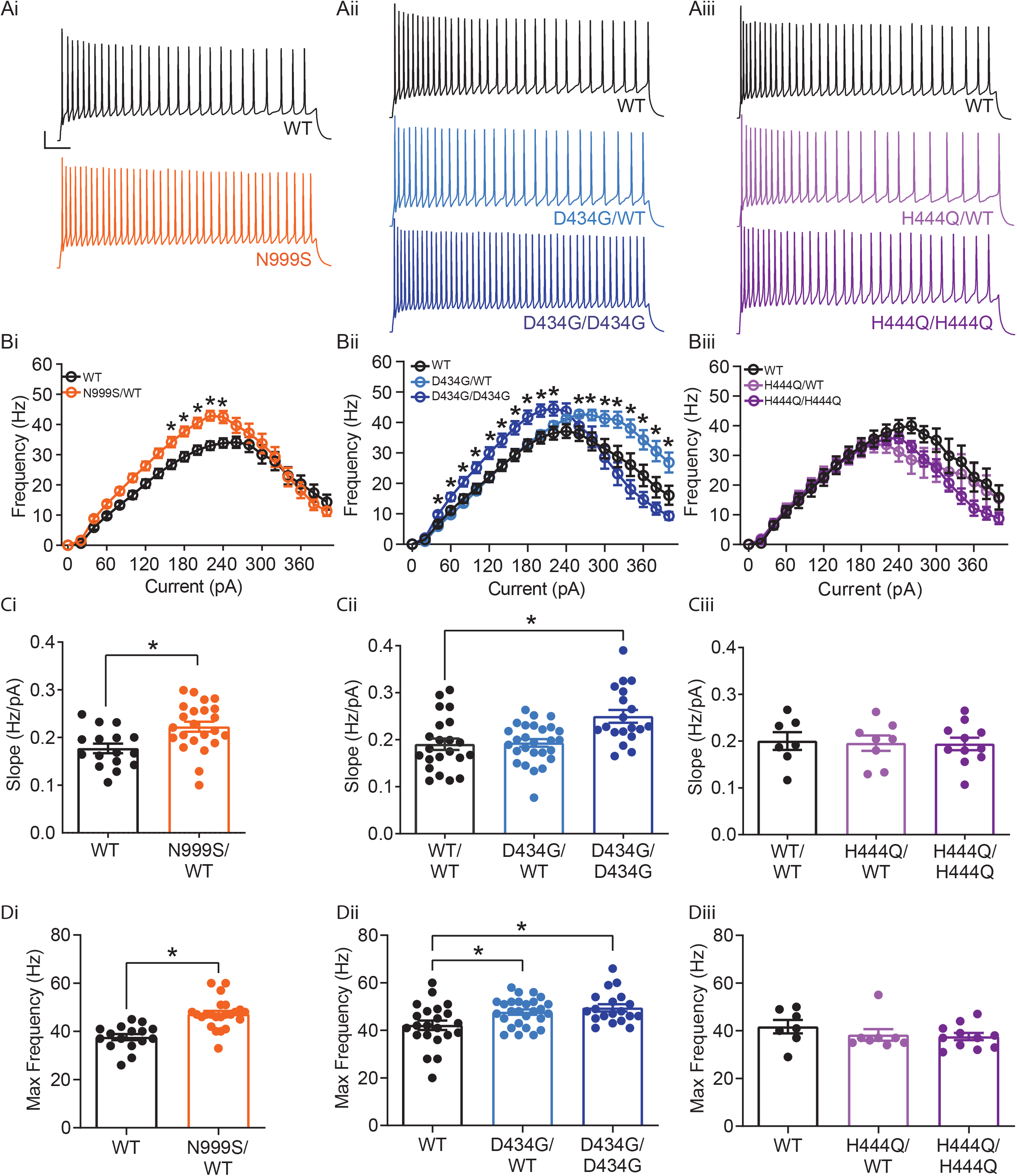
Increased intrinsic excitability in *Kcnma1*^N999S/WT^, *Kcnma1*^D434G/WT^, and *Kcnma1*^D434G/D434G^ granule neurons. In current-clamp mode, step currents from 0 to 400 pA were applied to dentate granule neurons under the same ionic conditions used to record BK currents. (A*i*-A*iii*) Representative AP trains elicited from the 200 pA current injection step in WT and transgenic neurons. Scale bar: 20 mV, 100 ms. (B*i*-B*iii*) Input-output relationship for firing frequency versus step current injection. Data are presented as mean ± SEM. *, *p* < 0.05, Two-way repeated measures ANOVA with Bonferroni post hoc. (B*i*) *Kcnma1*^N999S/WT^ (n=23 neurons, 5 mice) firing was higher than *Kcnma1*^WT/WT^ (n=16 neurons, 5 mice) at 160 pA (*p*=0.0426), 180 pA (*p*=0.0143), 200 (*p*=0.0068), 220 pA (*p*=0.0009), and 240 pA (*p*=0.0337) current steps. (B*ii*) *Kcnma1*^D434G/WT^ (n=27 neurons, 5 mice) firing was higher than *Kcnma1*^WT/WT^ (n=22 neurons, 5 mice) at 260 pA (*p=*0.0452), 280 (*p*=0.0314), 300 (*p*=0.0351), 320 (*p*=0.0177), 340 (*p*=0.0309), 360 (*p*=0.0358), 380 (*p*=0.0312), and 400 (*p*=0.0444) current steps. *Kcnma1*^D434G/D434G^ (n=19 neurons, 4 mice) firing was higher than *Kcnma1*^WT/WT^ at 40 pA (*p=*0.0266), 60 (*p*=0.0233), 80 (*p*=0.0277), 100 (*p*=0.0130), 120 (*p*=0.0074), 140 (*p*=0.0119), 160 (*p*=0.0084), 180 (*p*=0.0063), 200 (*p*=0.0059), and 220 (*p*=0.0261) current steps. (B*iii*) *Kcnma1*^H444Q/WT^ (n=8 neurons, 2 mice) and *Kcnma1*^H444Q/H444Q^ (n=11 neurons, 2 mice) firing was not different than *Kcnma1*^WT/WT^ (n=7 neurons, 1 mouse) at any current step (*p*=0.3222). (C*i*-C*iii*) Initial slope for the firing rate gain between 0 to 160 pA current injections. Data are presented as mean ± SEM, with individual data points. (C*i*) *Kcnma1*^N999S/WT^ firing slope was increased compared to WT (\**p*=0.0034; t-test). (C*ii*) *Kcnma1* ^D434G/D434G^ firing slope was increased compared to WT (\**p=*0.0051; One-way ANOVA), *Kcnma1* ^D434G/WT^ slopes were unchanged (*p*=0.9774). (C*iii*) *Kcnma1*^H444Q/WT^ and or *Kcnma1*^H444Q/H444Q^ firing slopes were not different than WT (*p*=0.9658). (D*i*) Maximal firing from *Kcnma1*^N999S/WT^ neurons was increased compared to WT (\**p*<0.0001; t-test). (D*ii*) Maximal firing from *Kcnma1* ^D434G/WT^ and *Kcnma1* ^D434G/D434G^ neurons was increased compared to WT (\**p*=0.0387 and *p*=0.0111, respectively; One-way ANOVA). (D*iii*) Maximal firing from *Kcnma1*^H444Q/WT^ and or *Kcnma1*^H444Q/H444Q^ neurons was not different than WT (*p*=0.4625; One- way ANOVA).

Increased firing was also observed in *Kcnma1*^D434G/WT^ neurons, but the shape of the input-output alteration was different than that observed in *Kcnma1*^N999S/WT^. Firing was 18-67% greater than WT controls at higher current injections only, from 260 to 400 pA (Figure 3A*ii*, B*ii*). Despite the increase at the higher end of the range, *Kcnma1*^D434G/WT^ firing still decreased after reaching a maximum, while remaining higher than *Kcnma1*^WT/WT^. The initial firing rate slope was not different from *Kcnma1*^WT/WT^ (Figure 3C*ii*). However, the maximal firing rate was 5.5 ± 2.2 Hz greater (113%) for *Kcnma1*^D434G/WT^ compared to *Kcnma1*^WT/WT^ (Figure 3D*ii*). This increase was shifted to higher current injections and occurred over a wider range of voltages than that observed for *Kcnma1*^N999S/WT^.

*Kcnma1*^D434G/D434G^ neurons, which had the highest BK current levels, were further different than *Kcnma1*^D434G/WT^. Firing was increased 22-47% in the early and middle of the current injection range, from 40 to 220 pA (Figure 3A*ii*, B*ii*). Both the initial slope (*Kcnma1*^D434G/D434G^ 0.25 ± 0.01 Hz/pA versus 0.19 ± 0.01 Hz/pA for *Kcnma1*^WT/WT^) and the maximal firing rate were greater (117%, Figure 3C*ii,* D*ii*). Yet *Kcnma1*^D434G/D434G^ firing was more qualitatively more similar to *Kcnma1*^N999S/WT^, despite the finding that *Kcnma1*^D434G/D434G^ BK current levels were almost twice as much as those recorded from *Kcnma1*^N999S/WT^.

No significant differences in firing frequency, slope for the initial firing rate gain, or maximal firing rate were observed in *Kcnma1*^H444Q/WT^ or *Kcnma1*^H444Q/H444Q^ neurons compared to WT littermates (Figure 3A-D*iii*). The lack of significant alteration in excitability was congruent with the absence of change in BK current levels in *Kcnma1*^H444Q/WT^ neurons.

We conclude that both the N999S and D434G GOF variants have pathogenic potential through their ability to increase BK currents and action potential firing. The LOF H444Q variant does not substantiate the same pathogenic potential under these conditions. Mechanistically, despite the grossly similar BK current levels between *Kcnma1*^D434G/WT^ and *Kcnma1*^N999S/WT^ neurons, the non-identical input-output curves suggest a more complex relationship between BK channel properties and neuronal excitability in dentate granule neurons. These mechanisms could manifest in either the passive membrane properties or at different phases of the action potential, related to the differential Ca^2+^ sensitivity for their GOF effects (Li *et al*., 2018; Moldenhauer *et al*., 2020; Yang, 2010). However, no additional differences in resting membrane potential (Vm) or input resistance (Ri) were observed between *Kcnma1*^N999S/WT^, *Kcnma1*^D434G/WT^, or *Kcnma1*^D434G/D434G^ and WT controls (Table S1). Analysis of AP waveforms from the 200 pA step suggested the basis for increased firing in *Kcnma1*^N999S/WT^ and *Kcnma1*^D434G/D434G^ neurons was a faster AHP decay (Figure S3), which would facilitate more rapid initiation of the next action potential. This AHP change was not significant in *Kcnma1*^D434G/WT^ neurons, consistent with the lack of alteration in firing at the same portion of the input-output curve.

### N999S and D434G reduce seizure thresholds in mice

Neuronal hyperexcitability is coincident with establishment of an epileptic network, and about half of all individuals with *KCNMA1* channelopathy, including those with N999S, D434G, and H444Q variants, report a history of seizures or epilepsy (Bailey *et al*., 2019; Miller *et al*., 2021). Individuals harboring the D434G variant primarily have absence seizures, if present (Du *et al*., 2005). Dentate gyrus hyperexcitability can both contribute to, and result from, epileptiform activity (Dengler *et al*., 2017; Krook-Magnuson *et al*., 2015; Mehranfard *et al*., 2015; Scharfman, 2019). In β4^—/—^ mice, increased granule neuron firing is found in the setting of hippocampal epileptiform discharges, non-convulsive seizures, and lower chemoconvulsant-induced seizure thresholds (Brenner *et al*., 2005; Whitmire *et al*., 2017). We hypothesized that *Kcnma1*^N999S/WT^ and *Kcnma1*^D434G/WT^ mice would show increased number, duration, or severity of seizure events compared to WT controls. However, since half of those harboring LOF variants also report seizures (Liang *et al*., 2019; Miller *et al*., 2021), including the H444Q and individuals with putative truncation alleles, *Kcnma1*^H444Q/WT^ and *Kcnma1*^—/—^ mice were assessed in parallel. No seizures have been previously reported in two established *Kcnma1*^—/—^ mouse models (Bailey *et al*., 2019; ALM unpublished data), but spontaneous epilepsy was reported in a CRISPR-exon4 *Kcnma1*^—/—^ line (Yao *et al*., 2021).

Behavioral assessments and EEGs were made from transgenic and WT littermates for indications of seizure. No spontaneous twitching/jumping/convulsions, rigidity/immobility, anorexia/dehydration, or premature mortality were observed from transgenic (or control) mice in the home-cage environment. After dural electrode implantation, 24-hr baseline EEGs were recorded. No interictal epileptiform discharges, spontaneous seizures, or other abnormalities (e.g., slowing) were observed in transgenic or control mice during baselines. The absence of spontaneous events was not surprising given that half of affected individuals do not report epilepsy, and among those that do, there is a wide range in frequency (isolated to daily), semiology, and age of onset (Bailey *et al*., 2019; Miller *et al*., 2021). However, this presents challenges to evaluating spontaneous EEG events in mouse models, especially those that could be occurring in deeper brain regions similar to β4^—/—^ mice. The presence of EEG abnormalities could be more comprehensively assessed with longer monitoring, depth electrodes, or interrogation of additional ages and strain backgrounds (Loscher *et al*., 2017), which were beyond the capability of the present study.

Human epilepsy variants in rodent models without spontaneous abnormalities often exhibit decreased thresholds to triggered seizures (Feliciano *et al*., 2011; Watanabe *et al*., 2000; Yuskaitis *et al*., 2018), although this is not entirely predictive of epilepsy risk (Noebels, 2003). We hypothesized that *Kcnma1*^N999S/WT^ and *Kcnma1*^D434G/WT^ mice would show either decreased threshold or increased severity with 40 mg/kg Pentylenetetrazol (PTZ) chemoconvulsant challenge. *Kcnma1*^WT/WT^ controls for each line developed seizures consistent with those observed with PTZ in other studies (Van Erum *et al*., 2019) ranging from abnormal posturing and myoclonic twitching (10/18 mice; modified Racine score 1 or 2) to tonic-clonic activity (7/18 mice; modified Racine 3 or 4) within minutes after PTZ injection (Video S1).

*Kcnma1*^N999S/WT^ mice developed PTZ-induced seizures that were distinguishable from *Kcnma1*^WT/WT^ littermates in several parameters. Behaviorally, most *Kcnma1*^N999S/WT^ mice displayed tonic-clonic activity (9/13 mice modified Racine 3 or 4), with two reaching status epilepticus (2/13 mice; modified Racine 5). The latency to first seizure after PTZ injection was reduced to 75 ± 15 sec, compared to WT littermates (294 ± 99 sec; Figure 4Ai, C-D). EEG power, an estimation of seizure severity, showed a broader range with *Kcnma1*^N999S/WT^ mice, although the differences were not significant (Figure 4Bi, C-D). Interestingly despite these observations, mice exhibiting electrographic seizures did not look strikingly behaviorally different from control mice. One reason may be the movement suppression that developed in *Kcnma1*^N999S/WT^ mice after PTZ injection, quantified by EMG. After PTZ, *Kcnma1*^WT/WT^ mice had infrequent bouts of sustained quiescent EMG activity, with average lengths of 45 ± 7 sec (n = 16). However, the inactive bouts were longer for *Kcnma1*^N999S/WT^ mice (311 ± 126, n= 10, P < 0.0001, Mann Whitney test) and were visually apparent (Video S2). The movement suppression exhibited by *Kcnma1*^N999S/WT^ mice under PTZ does not have a correlate in individuals harboring N999S variants, although a few report absence seizures among other types (Miller *et al*., 2021). Since no spontaneous EEG^+^/EMG*^—^* events were observed in the baseline EEG recording period of these mice, it remains to be determined whether the PTZ-elicited movement suppression is related to an absence-like seizure manifestation.

**Figure 4.**
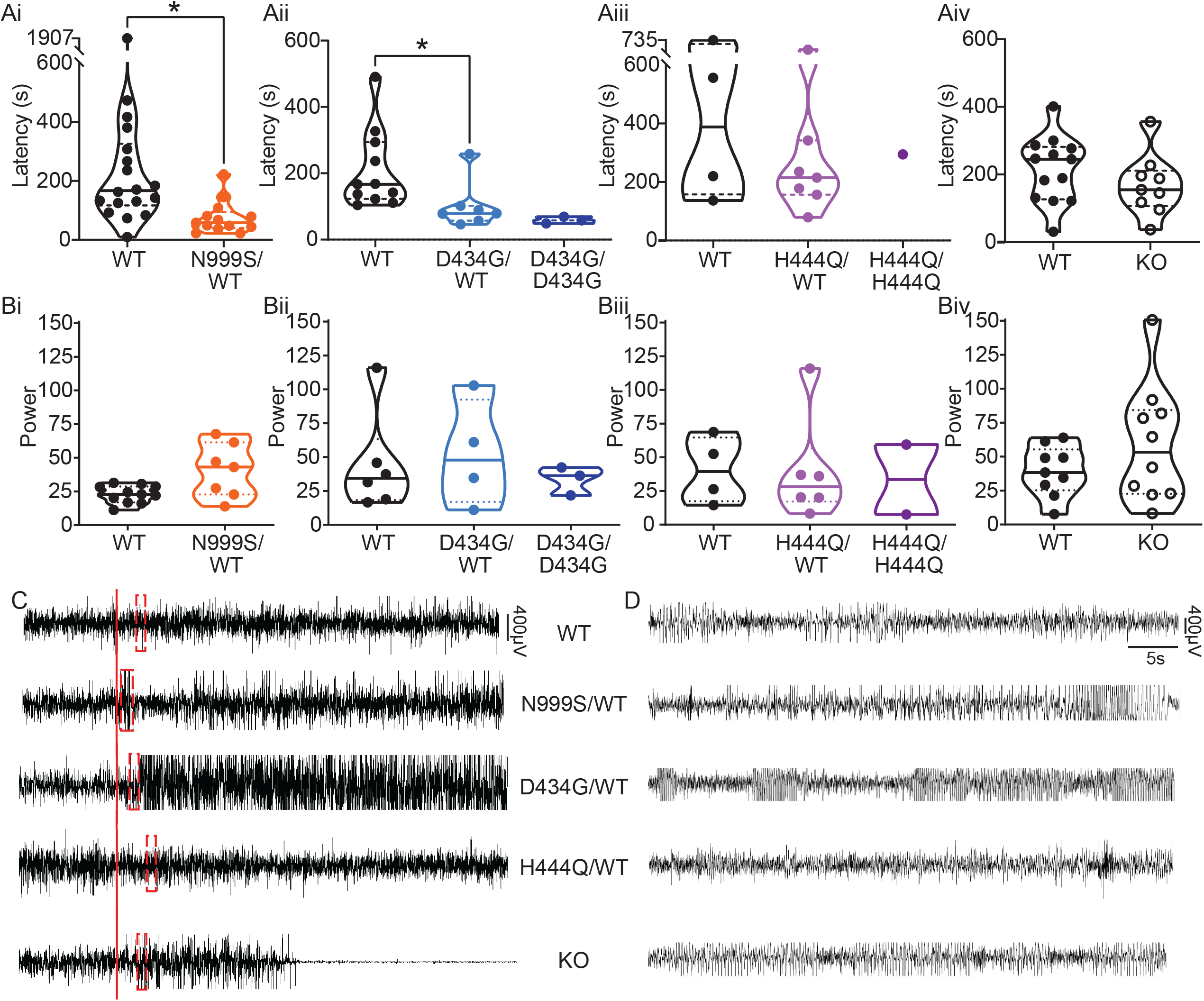
PTZ-induced seizures in mice. (A*i*-A*iv*) Latency to initial seizure after PTZ injection. Data are individual mice with median and inter-quartile range. (A*i*) Latency was decreased in *Kcnma1*^N999S/WT^ mice (n=13) compared to *Kcnma1*^WT/WT^ (n=18, \**p*=0.0006; Mann‒ Whitney test). (A*ii*) Latency was decreased in *Kcnma1*^D434G/WT^ mice (n=7) compared to *Kcnma1*^WT/WT^ (n=11, \**p*=0.0041; Mann‒Whitney test). *Kcnma1*^D434G/D434G^ mice were not included in the statistical analysis due to small sample size (n=3). (A*iii*) Seizure latency was comparable between *Kcnma1*^H444Q/WT^ (n=7) and *Kcnma1*^WT/WT^ (n=4, *p*=0.5273; Mann‒Whitney test). *Kcnma1*^H444Q/H444Q^ mice were not included in the statistical analysis due to small sample size (n=1). (A*iv*) No differences were found in seizure latency between *Kcnma1*^—/—^ (n=9) and *Kcnma1*^+/+^ mice (n=13, *p*=0.2282; Mann‒Whitney test). (B*i-iv)* Total EEG power after PTZ injection (y-axis in µV^2^/Hz x10^2^). Data are individual mice with median and inter-quartile range. (B*i*) EEG power was not different between *Kcnma1*^N999S/WT^ (n=7) and *Kcnma1*^WT/WT^ (n=11, *p*=0.0619; t-test). (B*ii*) *Kcnma1*^D434G/WT^ (n=4) was not different from *Kcnma1*^WT/WT^ (n=6, *p*=0.7563; t-test). (B*iii*) *Kcnma1*^H444Q/WT^ (n=6) was not different from *Kcnma1*^WT/WT^ (n=4, *p*=0.9641; t-test). (B*iv*) *Kcnma1*^—/—^ (n=10) was not different from *Kcnma1*^+/+^ (n=9, *p*=0.2134; t- test). (C) Representative EEG traces over 45 mins at baseline and after PTZ injection (red line). (D) Expanded EEG traces for the first seizure indicated with the red boxes in (C).

Within the D434G family, there an intermediate penetrance for epilepsy (56%), the most frequent diagnosis being absence (Du *et al*., 2005; Miller *et al*., 2021). Like N999S, *Kcnma1*^D434G/WT^ mice also showed a reduced latency to first seizure (101 ± 27 sec) compared to *Kcnma1*^WT/WT^ mice (209 ± 35 sec; Figure 4Aii, C-D). However, this reduction was not as large as the difference between *Kcnma1*^N999S/WT^ mice and their respective controls. Total EEG power from *Kcnma1*^D434G/WT^ mice was not different from WT controls (Figure 4Bii). Due to the low number of mice with EMG recordings (n = 3), no conclusions could be made regarding the presence of movement suppression after PTZ injection (Video S3). Therefore, the D434G variant also increased the propensity for seizure in the transgenic model, consistent with its ability to alter neuronal excitability, but was less severe than the N999S variant. The phenotype assessed here for *Kcnma1*^D434G/WT^ is also less severe than reported in a knock-in mouse model with the D434G mutation introduced in the context of a Cre/lox cassette. Those mice showed spontaneous spike-wave discharges in both the heterozygous and homozygous configuration with complete penetrance (Dong *et al*., 2021), a phenotype that appears more severe than reported in the D434G pedigree, in which only half experience seizures (Du *et al*., 2005).

Although no patients have a homozygous D434G genotype, a limited number of *Kcnma1*^D434G/D434G^ mice were available for EEG analysis. We tested whether *Kcnma1*^D434G/D434G^ mice, producing only mutant BK channel homotetramers, had a more severe phenotype. These mice showed a trend toward the shortest latencies to seizure, with thresholds comparable to the lowest among the *Kcnma1*^D434G/WT^ cohort (Figure 4Aii, C-D). *Kcnma1*^D434G/D434G^ mice had EEG power overlapping with WT and heterozygous littermates (Figure 4Bii). While not conclusive, the data are suggestive of a gene effect with D434G dosage that further reduces seizure threshold.

For LOF transgenics, changes in seizure threshold in both directions were considered. Approximately the same proportion of individuals with LOF variants report seizure as those with GOF variants (Miller *et al*., 2021), a finding validated in one *Kcnma1*^—/—^ mouse model (Yao *et al*., 2021). Yet paradoxically, acute inhibition of BK channels has anti-seizure effects in other rodent models (Dong *et al*., 2021; Kuebler *et al*., 2001; Sheehan *et al*., 2009). However, no differences were observed in latency to first seizure or total EEG power in *Kcnma1*^H444Q/WT^ or *Kcnma1*^—/—^ mice (Figure 4Aiii-Aiv, Biii-Biv, C-D; Video S4). For H444Q, this data suggests the lack of change in dentate granule neuron BK currents and excitability may be consistent with other areas of the brain, producing no change in seizure propensity indicative of widespread hyperexcitability. Overall, alterations in BK current and firing detected concurrently with lowered seizure threshold suggests the GOF variants N999S and D434G have the potential to contribute to seizure risk by changing neuronal activity in a mouse model. However, partial (H444Q) or total (*Kcnma1*^—/—^) loss of BK channel function does not support the same potential in seizure etiology under equivalent conditions.

### N999S and D434G cause paroxysmal dyskinesia in mice

One of the most recognizable symptoms in *KCNMA1* channelopathy is a distinctive type of dyskinesia manifesting as sudden, brief paroxysms of axial hypotonia with dystonia (PNKD3). These episodes sometimes resemble the immobility in non-narcoleptic cataplexy, but have preservation of some muscle tone that varies among individuals (Du *et al*., 2005; Heim *et al*., 2020; Keros *et al*., 2022; Miller *et al*., 2021; Wang *et al*., 2017; Zhang *et al*., 2015). Patients may slump or fall over but can often maintain position if appropriately supported, and consciousness is maintained. Normal activity is recovered relatively quickly without persistent impairment (see patient videos in Braverman, 2019; Sanders, 2018). PNKD3 episodes are not initiated by movement or exertion (non-kinesigenic), but rather by triggers such as stress, strong emotion, cold, fatigue, or alcohol. The events are not associated with epileptiform activity on EEG and are generally unresponsive to anti-seizure medications (Keros *et al*., 2022; Miller *et al*., 2021). PNKD3 is associated with substantial morbidity due to its high frequency, with hundreds of episodes per day. All three variants tested in this study are associated with PNKD, with 75% of individuals harboring N999S and D434G carrying the diagnosis but also observed at lower incidence with LOF variants or VUS (Miller *et al*., 2021).

There are currently no standardized behavioral assays for either PNKD3 or non- *KCNMA1*-associated PNKD. In other paroxysmal dyskinesia animal models, the phenotype is usually hyperkinetic, not the hypotonic events observed in PNKD3. For example, Ca^2+^ channelopathy, *PRRT2-*deficient, and *PNKD* mutant mice are characterized by dystonia, chorea, and tonic-clonic episodes (Fureman *et al*., 2002; Lee *et al*., 2012; Michetti *et al*., 2017; Tan *et al*., 2018). No spontaneous hypotonic dyskinetic motor behavior was detectable to a blinded observer in any of the transgenic lines in this study. Therefore, we sought to elicit episodes by utilizing known triggers for PNKD3. Since individuals harboring N999S and H444Q variants are mostly children without any reported alcohol exposures, a PNKD trigger specifically reported for D434G (Du *et al*., 2005), and no calibratable emotional responses are validated in mice, we focused on the standardizable stress experienced during physical restraint. Stress provocation is the closest stimulus to the natural triggers observed in PNKD3-affected individuals (Miller *et al*., 2021). Restraint stress provoked dyskinesia in most (85%) *tottering* mice (Fureman *et al*., 2002), and *PNKD* mutant mice also showed dyskinesia after stressful handling when placed in a beaker (Lee *et al*., 2012).

To test whether restraint stress would produce paroxysmal dyskinesia, mice were subjected to an acute stereotypical manual restraint protocol by an experienced handler. After restraint, mice were placed into a beaker, a novel constrained environment proposed to enhance stress (Lee *et al*., 2012). Mice with PNKD-like characteristics were predicted to show restraint- triggered episodes of hypotonia. Mice were scored for abnormal movement (time immobile, circling/hyperactivity, twisting/chorea or limb-clasping, tonic-clonic movement, flattened/dystonic posturing, tremor, listing and falling) in the beaker under video observation. Stereotypical behaviors such as grooming were also recorded. WT mice from all groups showed normal exploratory behavior including sniffing, grooming and rearing with coordinated movements.

*Kcnma1*^N999S/WT^ mice and WT littermates placed in the beaker without prior restraint did not show any dyskinetic movements or collapsing behavior (paws no longer touching the ground). There was no significant difference in the time spent immobile between these groups (Figure 5A). Next, restrained mice were placed in the beaker. Since mice increase grooming when released from stress (Crawley, 2000), this behavior was used as a control to indicate the presence of stress. *Kcnma1*^N999S/WT^ mice and WT littermates both showed an increase in grooming events after restraint compared to their non-restraint controls (Figure 5B), confirming both genotypes responded to stress with an increase in stereotypical behavior.

**Figure 5.**
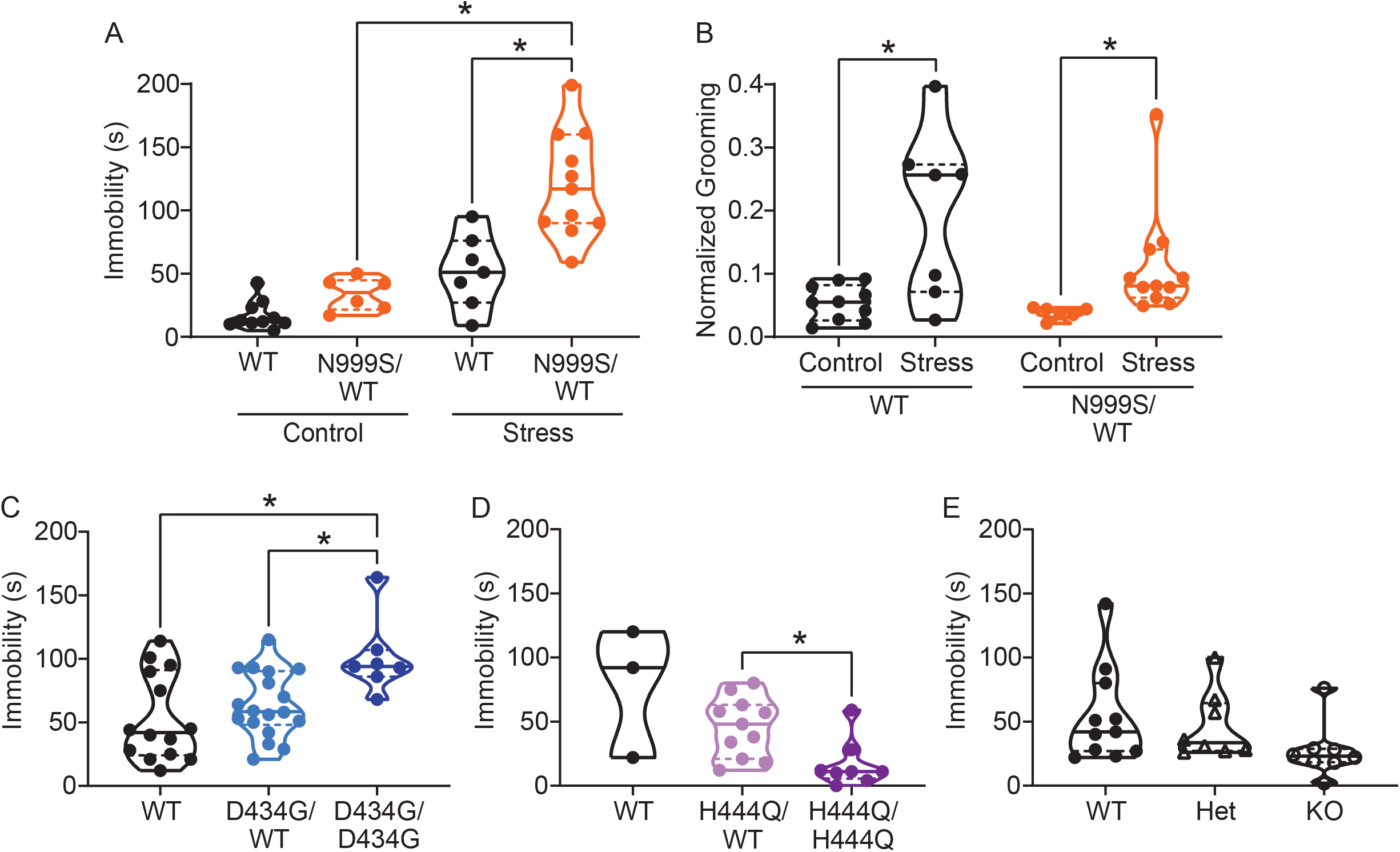
Stress-induced paroxysmal dyskinesia. (A) Control: Without restraint stress, there was no difference in the time spent immobile between *Kcnma1*^WT/WT^ (n=10) and *Kcnma1*^N999S/WT^ mice (n=6, *p*>0.9999; Two-way ANOVA with Bonferroni posthoc). Restraint stress: Immobility time was longer for restrained *Kcnma1*^N999S/WT^ mice (n=11) compared to *Kcnma1*^WT/WT^ (n=7, \**p*=0.0001; One-way ANOVA), and between restrained *Kcnma1*^N999S/WT^ mice (n=11) compared to unrestrained *Kcnma1*^N999S/WT^ mice (n=6, \**p*<0.0001). In contrast, unrestrained *Kcnma1*^WT/WT^ mice (n=10) had no differences from restrained *Kcnma1*^WT/WT^ mice (n=7, *p*=0.1174). (B) Grooming behavior increased in restrained *Kcnma1*^WT/WT^ mice (n=7) compared to unrestrained *Kcnma1*^WT/WT^ mice (n=10, \**p*=0.0300; t-test), and in restrained *Kcnma1*^N999S/WT^ mice (n=11) compared to unrestrained *Kcnma1*^N999S/WT^ mice (n=6, \**p=*0.0174; t-test). (C) After restraint, *Kcnma1*^D434G/D434G^ mice (n=7) spent more time immobile compared to *Kcnma1*^WT/WT^ mice (n=14, \**p*=0.0166; One-way ANOVA). However, *Kcnma1*^D434G/WT^ mice were not different (n=18, *p*=0.7174). (D) Immobility time was shorter in restrained *Kcnma1*^H444Q/H444Q^ mice (n=8) compared to *Kcnma1*^H444Q/WT^ mice (n=11, \**p*=0.0081; t-test). *Kcnma1*^WT/WT^ mice were not included in the statistical analysis due to small sample size (n=3). (E) *Kcnma1*^−/−^ mice (n=8) had reduced immobility compared to *Kcnma1*^−/+^ mice (n=8) and *Kcnma1*^+/+^ mice (n=11, *p*=0.0535; Kruskal-Wallis test). Data are individual mice with median and inter-quartile range.

After stress, *Kcnma1*^WT/WT^ mice had exploratory behavior and spent less than a minute immobile in the beaker (51 ± 10 sec). Although the range was wider, their time spent immobile did not differ significantly from the unrestrained baseline. In contrast, *Kcnma1*^N999S/WT^ mice were immobile for more than twice as long after stress (120 ± 12 s)(Figure 5A, Video S5). After episodes of immobility, both genotypes resumed normal exploratory behavior or grooming.

In qualitative assessment, three *Kcnma1*^N999S/WT^ mice had extended myoclonic ‘hiccups’ throughout the immobility that were not associated with respiratory rate. One mouse also showed listing, and three had a flattened posture. Evaluation of other dyskinetic behaviors (dystonia, chorea, clasping, etc) in non-restraint controls and after stress revealed grossly normal movements for *Kcnma1*^N999S/WT^ mice, with the exception of the notable immobility. In direct comparison, *Kcnma1*^WT/WT^ littermates had raised heads and less hunched postures during their briefer immobility, suggesting the maintenance of normal axial tone. Brief hiccups were observed in one WT control, at shorter duration than the *Kcnma1*^N999S/WT^ mice, and one mouse had a brief flattened posture during the first bin. Use of a fitted tube restraint, which may produce a stronger stress response, increased the ability of a blinded observer to predict genotype differences in immobility (*n* = 11 mice, data not shown).

Taken together, these data suggest that the presence of a stressor (restraint) produces a new behavioral state in *Kcnma1*^N999S/WT^ mice (immobility) that was not observed in the absence of the trigger or in WT littermates. If the immobility behavior resulted from stress-induced atonic or absence seizures, these events would likely have been observed during baseline EEG recordings given the number of occurrences in the five-minute beaker assay. If stress-induced immobility resulted from general hypoactivity or altered fear response, open field testing might show a difference in motor exploratory behavior between *Kcnma1*^N999S/WT^ and WT littermates, which was not observed (Figure S4). *Kcnma1*^N999S/WT^ mice were also able to achieve the same peak speed as WT littermates during voluntary wheel running (Figure S5Bi). In addition, there is no evidence from patients for correlation of PNKD3 with increased anxiety, depression, or hypoactivity (Miller *et al*., 2021). We conclude that stress-induced immobility, which occurs in brief episodes that are instantaneously recovered, and without other hyperkinetic or tonic-clonic manifestations, is consistent with the reversible triggered hypotonic behavioral state in PNKD3- affected individuals (Heim *et al*., 2020; Keros *et al*., 2022).

Besides stress, PNKD3 episodes can be triggered by positive emotions or excitement, similar cataplexy in patients with narcolepsy (Dauvilliers *et al*., 2014; Miller *et al*., 2021; Sun *et al*., 2019). Related to the reward and arousal effects in mice, cataplexy can be provoked in narcoleptic orexin^—/—^ mice by wheel running (Espana *et al*., 2007; Mahoney *et al*., 2017; Novak *et al*., 2012). We also assessed this positive trigger to determine if voluntary wheel running could produce a PNKD-like behavior in the setting of a more complex motor task. In this assay, *Kcnma1*^N999S/WT^ mice covered a shorter distance compared to their WT littermates, and they had an increase in the number of gaps between running (Figure S5Ai, Di). This phenotype is consistent with the presence of, but not exclusively attributable to, PNKD since *Kcnma1*^N999S/WT^ mice also show baseline motor dysfunction in the rotarod and hanging wire assays (Figure S6A, E).

PNKD3 is also exhibited in individuals harboring heterozygous D434G variants and is provoked by additional triggers besides stress, such as alcohol (Du *et al*., 2005). However, without diagnostic standardization, it is not clear whether this constitutes a different type of PNKD episode in individuals harboring D434G compared to N999S. In the stress assay, *Kcnma1*^D434G/WT^ mice and WT littermates had similar immobility lasting 53 ± 9 sec and 63 ± 6 sec, respectively (Figure 5C). However, in homozygous *Kcnma1*^D434G/D434G^ mice, immobility time was similar to N999S heterozygotes (101 ± 11 sec). Thus, stress-induced dyskinesia is present in the D434G mouse model, and the increasing allelic severity was consistent with both seizure assays and the degree of BK channel impairment in heterologous cells. It remains possible that alcohol would also be capable of triggering these episodes, but it is difficult to assess given the bi-directional motor effects of alcohol in mice (Crawley, 2000).

In contrast, homozygous LOF manipulations showed a different directionality in stress-triggered dyskinetic behavior, with less immobility after restraint than either WT or heterozygous littermates. No phenotypic differences were detected in heterozygous mice of either line. However, immobility in *Kcnma1*^H444Q/H444Q^ mice was reduced to 16 ± 6 sec (Figure 5D), and in *Kcnma1*^—/—^ mice, it was reduced to 27 ± 7 sec (Figure 5E). Three *Kcnma1*^—/—^ mice also showed hyperactive circling and rapid limb movements notable to the blinded observer, and one had notable tremor during the brief non-active periods. These data reveal that the H444Q variant and *KCNMA1* null genotypes are not associated with PNKD-immobility under the same triggers that provoke GOF variants. The results raise the possibility that stress-induced dyskinesia manifestation is influenced by mutation type, with GOF producing hypokinetic and LOF producing hyperkinetic responses.

## DISCUSSION

We have characterized the channel properties, neuronal activity, neurobehavioral phenotypes, and relative severity of three *KCNMA1*–linked channelopathy variants under equivalent conditions. Pathogenic potential was established using four criteria: 1) low variant frequency in the human population, classifying as a mutation (Miller *et al*., 2021), 2) variant alters BK channel gating properties, 3) variant alters neuronal BK currents and firing, since the channelopathy is a neurological disorder, and 4) variant produces phenotypes similar to the central patient diagnoses, seizure susceptibility and PNKD. The findings support the conclusion that *KCNMA1*-linked channelopathy, although symptomatically heterogenous and comprised predominantly of *de novo* variants, has the potential to be categorized as a monogenic disorder. The results substantiate hyperexcitability, increased seizure propensity, and PNKD as collective phenotypes replicated in two hypermorphic GOF *KCNMA1* alleles. Moreover, our data for N999S and D434G corroborate mounting evidence, both in patients (Du *et al*., 2005; Keros *et al*., 2022; Miller *et al*., 2021; Zhang *et al*., 2020) and animal models (Dong *et al*., 2021; Kratschmer *et al*., 2021), that PNKD can be considered the most consistent symptom for *KCNMA1* GOF channelopathy.

It is not yet clear whether all *KCNMA1* variants discovered in the setting of neurological diagnoses carry the same pathogenic potential. *KCNMA1* LOF channelopathy has been proposed to carry a broader set of non-overlapping features associated with a subset of *de novo* LOF variants, referred to as Liang-Wang syndrome (Liang *et al*., 2019). However, none of the observable patient correlates were present in the LOF H444Q model studied here. H444Q mice also had little overlap with *Ryegrass Staggers*, a toxicity syndrome of livestock involving BK channel inhibition that is phenotypically similar to *Kcnma1*^—/—^ mice (Imlach *et al*., 2008).

Although N999S is the most commonly reported *KCNMA1* variant, direct evidence that it caused channelopathy was lacking because it arose *de novo* in all known cases (Keros *et al*., 2022; Miller *et al*., 2021). In mice, our data validate its dominant inheritance and pathogenic potential as a GOF mutation able to increase BK current and neuronal activity in the heterozygous configuration in mice. Neurobehavioral validation further identified increased PTZ-induced seizure propensity and stress-triggered dyskinesia episodes resembling PNKD-like immobility, broadly consistent with the phenotypic occurrence in patients. The lethality of homozygous *Kcnma1*^N999S/N999S^ (and hemizygous *Kcnma1*^N999S/Δ^) genotypes, which have not been found in any patient, underscore the severity of this variant.

D434G, a less severe GOF mutation at the BK channel level, had dominant inheritance for a subset of traits in our model, partially paralleling the familial pedigree (Du *et al*., 2005). The increased BK current, excitability, and PTZ-induced seizure propensity in the heterozygous configuration validated D434G pathogenicity. However, homozygosity was required to produce the PNKD-like immobility attacks, ranking D434G less pathogenic than N999S. Interestingly, three other D434G models have been reported, with some phenotypic variation. BAC-loxP- D434G mice exhibited generalized tonic-clonic seizures in the absence of motor dysfunction (Ling, 2016). Another Cre/lox-D434G mouse line was comparatively more severe with complete penetrance of absence seizures and dyskinesia in the heterozygous configuration (Dong *et al*., 2021). This variability, observed both in mouse models and incomplete penetrance in patients, raises the possibility that additional genetic or environmental factors can influence symptomatic severity. Nevertheless, an analogous mutation in flies also alters neuronal activity and baseline motor behavior (Kratschmer *et al*., 2021).

The LOF variant, H444Q, demonstrated limited pathogenicity by decreasing BK channel activity but was not validated as a hypomorphic or haploinsufficient allele in neurons or mice. Homozygous *Kcnma1*^H444Q/H444Q^ mice showed a neurobehavioral phenotype distinct from PNKD: stress-induced hyperkinetic motor responses similar to *Kcnma1*^—/—^ null mice. Given multiple genetic findings and symptomatic ambiguity in the patient carrying this variant, the different dyskinetic responses compared to the two GOF models may suggest a basis to improve the diagnostic investigations for this and other *KCNMA1* variants classified as LOF or VUS. At present, there are no patients with homozygous *KCNMA1* alleles validated as functionally null for channel activity (Miller *et al*., 2021), but the ataxia, tremor, decreased strength, and hyperactivity in *Kcnma1*^—/—^ mice (Imlach *et al*., 2008; Meredith *et al*., 2004; Sausbier *et al*., 2004; Typlt *et al*., 2013; Wang *et al*., 2020) are symptoms observed at lower incidence among patients. Lastly, our experimental conditions failed to corroborate the influence of LOF alleles on seizure propensity predicted from several animal studies (Ermolinsky *et al*., 2008; Kuebler *et al*., 2001; Pacheco Otalora *et al*., 2008; Sheehan *et al*., 2009; Shruti *et al*., 2008; Yao *et al*., 2021).

Genotype-phenotype relationships are important for understanding *KCNMA1* channelopathy disease mechanisms as well as potential therapeutics. The allelic series established from BK channels *in vitro* is outwardly congruent with the relative severity in mice. Such allelic series have been pivotal in understanding other complex channelopathies, especially those delineating distinct disorders within the same gene association (Noebels, 2003; Pietrobon, 2005; Zwingman *et al*., 2001). However, because this *KCNMA1* allelic series was derived from a limited set of conditions, designations of phenotypic severity could be influenced by additional factors. Variant effects on channel expression were not addressed in this study, nor was their context within the mouse gene, which was not humanized through additional rounds of gene editing. Mouse and human BK channels differ at eight constitutive coding residues and have minor differences in BK current properties in heterologous cells (Lai, 2015). However, D434G produces a larger GOF effect on the G-V relationship in the context of a human BK channel compared to mouse (Wang *et al*., 2009). It is also not clear which BK channel properties or which neurons/circuits are the most critical for symptomology. BK current levels were relatively similar between *Kcnma1*^N999S/WT^ and *Kcnma1*^D434G/WT^ in one type of neuron (dentate granule), yet these two genotypes exhibited different motor behavior. Instead, homozygous *Kcnma1*^D434G/D434G^ PNKD was comparable to *Kcnma1*^N999S/WT^. Additional experiments to probe the voltage and Ca^2+^ dependent bases for N999S and D434G gating defects in neurons may further explain the relative pathogenicity (Diez-Sampedro *et al*., 2006; Du *et al*., 2005; Li *et al*., 2018; Moldenhauer *et al*., 2020; Wang *et al*., 2009; Yang, 2010).

Changes in BK channel function and/or *KCNMA1* expression are associated with a growing number of neurodevelopmental disorders including epilepsy, dyskinesia, autism, Angelman’s syndrome, Fragile X syndrome, and brain and skeletal malformations (Cheng *et al*., 2021; Deng and Klyachko, 2016; Du *et al*., 2020; Kessi *et al*., 2020; Laumonnier *et al*., 2006; Liang *et al*., 2019; Miller *et al*., 2021; N’Gouemo, 2014; Sun *et al*., 2019). Neuropathology in these disorders is associated with changes in BK channel activity in both directions. Yet it has been challenging to distill *KCNMA1*-linked channelopathy into a cohesive GOF versus LOF symptomology because the existing patient data lack genetic pedigrees and diagnostic cross-comparability.

Looking ahead, the phenotypic penetrance and heterogeneity investigated here validate only a few of the 40+ patient-associated *KCNMA1* variants, but it will not be possible to make transgenic models for every case. There is less symptomatic consistency among non-GOF alleles (LOF or VUS), identifying this as a potentially fruitful area for future investigations. Our mouse models may also provide insight into therapeutic mechanisms. Stimulants (Lisdexamfetamine and dextroamphetamine) are highly effective in reducing PNKD in multiple children harboring N999S variants, as well as the GOF variant N536H (Keros *et al*., 2022; Zhang *et al*., 2020). In heterologous cells, neither drug has a direct effect on BK^WT^ channel activity (Figure S1D and Zhang *et al*., 2020) or BK^N999S^ (Figure S1D), leaving the target for their actions on PNKD3 an open question for future mechanistic studies using these mice.

## MATERIALS AND METHODS

### HEK cell patch-clamp electrophysiology

The N999S (rs886039469; also numbered as N995S, N1036S, and N1053S in other reference sequences), D434G(rs137853333), and H444Q mutations were introduced into wild-type (WT) human hBKQEERL cDNA sequence (MG279689) in the pcDNA3.1+ mammalian expression vector. Mutations were verified by sequencing. Channel constructs contained an N-terminal Myc tag and an EYFP tag in RCK2 domain.

HEK293T cells (CRL-11268, ATCC, Manassas, VA, USA) were cultured in media containing Dulbecco’s modified Eagle medium (cat. #11995-065, Gibco, Life Technologies Corp., Grand Island, NY, USA), 10% fetal bovine serum (cat. #4135, Sigma-Aldrich, St. Louis, MO, USA), 1% penicillin/streptomycin (cat. #30-002-Cl, Mediatech Inc., Manassas, VA, USA), and 1% L-glutamine (cat. #25-005-Cl, Mediatech Inc., Manassas,VA, USA) and incubated with 5% carbon dioxide at 37°C. HEK cells were transfected with WT or mutant constructs using Trans-IT LT1 (Mirius Biological, Madison, WI, USA) at 1:2 ratio of DNA and the reagent. After 4–12 h, cells were re-plated onto glass coverslips pre-coated with poly-L-lysine (cat. #P4832, Sigma-Aldrich, St. Louis, MO, USA). After 14-24h, recordings were performed.

BK currents were recorded using inside-out patch-clamp at room temperature in symmetrical K^+^. 1 μM intracellular Ca^2+^ was used, a physiologically relevant Ca^2+^ condition near concentrations where altered gating behavior was manifested in prior studies (Li *et al*., 2018; Moldenhauer, 2020; Moldenhauer *et al*., 2020; Wang *et al*., 2009; Yang *et al*., 2010). Thin-walled borosilicate glass pipettes with resistances of 1–3 MΩ were filled with (in mM): 140 KMeSO3, 2 KCl, 2 MgCl2, and 20 HEPES. The internal (bath) solution contained (in mM): 140 KMeSO3, 2 KCl, 20 HEPES, and 5 HEDTA with CaCl2 added to achieve 1 μM free CaCl2, pH adjusted to 7.2 with KOH). Free Ca^2+^ concentrations were calculated with WebMaxC: (https://somapp.ucdmc.ucdavis.edu/pharmacology/bers/maxchelator/webmaxc/webmaxcS.htm).

Macroscopic currents were recorded with a Multiclamp 700B amplifier, and signals were filtered at 10 kHz and digitized at 50kHz using Digidata1440A and pCLAMP v10 software (Molecular Devices, Sunnyvale, CA, USA). BK currents were activated with a voltage protocol stepped from a holding potential of −100 mV stepping to +250 mV with +10 mV increments for 30 ms and back to −100 mV for 15 ms to generate tail currents. Conductance - voltage (G-V) curves were generated from the tail currents 150–200 µs after the peak normalized to the maximum conductance (G/Gmax) and plotted against the activating voltage step (V). V1/2 values were calculated from a Boltzmann fit of the G-V curves (Prism v9 GraphPad Software, San Diego, CA). Leak currents were compensated using a P/5 protocol with a subsweep holding potential of −120 mV.

### Generation of Kcnma1^N999S^, Kcnma1^D434G^, and Kcnma1^H444Q^ mouse lines

Heterozygous founders introducing N999S (AAT→ A**<u>G</u>**T, exon 25), D434G (GAT→ G**<u>G</u>**T, exon 10), and H444Q (CAC → CA**<u>G</u>**, exon 10) mutations into the mouse *Kcnma1* gene (Gene ID:16531) were generated with CRISPR/Cas9 homologous recombination methods in the C57BL/6J strain (Figure S1). *Kcnma1*^D434G/WT^ and *Kcnma1*^H444Q/WT^ mice were generated at the Transgenic Mouse Core at John Hopkins University (Baltimore, MD). *Kcnma1*^N999S/WT^ mice were generated at The Jackson Laboratory (Bar Harbor, ME). Transgenic mice were validated with *Kcnma1* sequencing, and founders without additional non-synonymous mutations were bred with C57BL/6J for N1 progeny at The University of Maryland School of Medicine. Genotyping was performed at Transnetyx, Inc (Cordova, TN) using primers described in Supplemental Methods. Transgenic lines were backcrossed with C57BL/6J up to four generations (N4) for experimental cohorts with heterozygous progeny.

N2-N4 heterozygous mice were intercrossed to produce homozygous progeny. Transgenic heterozygous *Kcnma1*^N999S/WT^, *Kcnma1*^D434G/WT^, *Kcnma1*^H444Q/WT^ (as well as homozygous *Kcnma1*^D434G/ D434G^ and *Kcnma1*^H444Q/ H444Q^) mice showed no gross differences in home cage behavior, body weights or gross morphology, or breeding. *Kcnma1*^N999S/WT^ x *Kcnma1*^N999S/WT^ heterozygous crosses produced either no viable pups (n = 5/10 breedings), or no homozygous pups (n = 56 pups). Additional crosses with *Kcnma1^—/+^* heterozygous dams mated to *Kcnma1*^N999S/WT^ males (n = 15 viable pups from 5 breedings) also produced no *Kcnma1*^N999S/*—*^pups.

*Kcnma1*^—/—^ (Slo KO) mice were maintained on a C57BL/6J background (> N12 generation). Littermates of each genotype were produced from heterozygous *Kcnma1*^+/—^ x *Kcnma1*^+/—^ breeding pairs, as described previously (Meredith *et al*., 2004) using primer sequences in Supplemental Methods. For all lines, male and female mice were separated by sex and group housed on a 12h light/12h dark cycle. Littermates from both sexes were used for experimental procedures, at the indicated ages. Chow and water were provided *ad libitum*.

*Kcnma1*^N999S/WT^ breeders were fed with high fat chow supplement to enhance breeding. All experiments were conducted in accordance with the University of Maryland School of Medicine Animal Care and Use Guidelines and approved by the Institutional Animal Care and Use Committee.

### Patient data

Patient phenotype and genetics data cited as ‘data not shown’ was exempt under University of Maryland School of Medicine Institutional Review Board (IRB) Non-Human Subjects Research (NHSR) Protocols HP-00083221, HP-00086440, and HP-00092434.

### Gene expression

Unilateral hippocampus and the medial portion of cerebellum were extracted from 4- month-old mice, and directly put them in 1.0 mm diameter zirconium beads with 750µl of trizol for bead homogenization. RNA was extracted using the miRNeasy Mini Kit (Qiagen, Valencia, CA) following manufacturer’s protocol. To assess the RNA quality, RNA was quantified via RNA-40 nanodrop and OD 260/280 ratio of all samples were between 1.94 and 2.05. Integrity of RNA was examined via Agilent 2,100 Bioanalyzer (Agilent Technologies, Palo Alto, CA, USA). RNA integrity number (RIN) of all samples were >9. For transcriptome analysis, mouse Clariom™ D Assay (Applied Biosystems™, Waltham, Massachusetts, USA.) was used following manufacturer’s instruction. RNA extraction and array processing were done at the Genomics Core Facility, University of Maryland, Baltimore. The raw microarray profiling data was preprocessed and quartile-normalized using the Transcriptome Analysis Console Software (Version 4.0.1) (https://www.thermofisher.com/us/en/home/life-science/microarray-analysis/microarray-analysis-instruments-software-services/microarray-analysis-software/affymetrix-transcriptome-analysis-console-software.html, accessed on 1 December 2021). All samples passed array quality control evaluation. Data normalization, differential expression, and hierarchical clustering analysis was performed with default parameters.

### Hippocampal Slice electrophysiology

Three to four-week-old mice were anesthetized with isoflurane, and brains were removed and placed into ice-cold sucrose dissection solution (in mM): 10 MgCl2, 26 NaHCO3, 1.25 Na2HPO4, 3.5 KCl, 0.05 CaCl2, 10 glucose, 200 sucrose, 1.2 sodium pyruvate, and 0.4 vitamin C, bubbled with 95% O2 and 5% CO2. The brain slices were cut coronally at 300 μm on a VT1000S vibratome (Leica Microsystems, Wetzlar, Germany) at 3–4 °C. Slices containing hippocampus were incubated at 32°C for 30 min and kept at the room temperature in oxygenated artificial cerebrospinal fluid (ACSF) containing (in mM): 125 NaCl, 1.7 MgCl2, 26 NaHCO3, 1.25 Na2HPO4, 3.5 KCl, 2 CaCl2, 10 glucose, 1.2 sodium pyruvate, and 0.4 vitamin C.

Whole cell patch clamp recordings were performed in the dentate gyrus granule cells (GCs) of the hippocampus using borosilicate glass electrodes pulled at 3-5 MΩ filled with (in mM): 123 K-MeSO3, 9 NaCl, 0.9 EGTA, 9 HEPES, 14 Tris-phosphocreatine, 2 Mg-ATP, 0.3 Tris-GTP, and 2 Na2-ATP, pH adjusted to 7.3 with KOH, 290-300 mOsm. The low EGTA intracellular solution allows endogenous Ca^2+^ influx to activate BK channels (Fakler and Adelman, 2008; Jackson *et al*., 2004; Muller *et al*., 2007; Whitt *et al*., 2018). The slices were perfused with oxygenated ACSF at room temperature during the recordings. Granule cells in the DG were visualized with a Luca-R DL-604 EMCCD camera (Andor, Belfast, UK) under IR-DIC illumination on an FN1 upright microscope (Nikon, Melville, NY, USA). Current- and voltage-clamp recordings were made with a Multiclamp 700B amplifier, and signals were filtered at 10 kHz and digitized at 50kHz using Digidata1440A and pCLAMP v10 software (Molecular Devices, Sunnyvale, CA, USA). All data were corrected for liquid junctional potential (10mV).

In current clamp recordings, the resting membrane potential was measured without any current input after a whole-cell mode was made. If the initial resting membrane potential was more depolarized than ‒70 mV or a series resistance was larger than 20MΩ, the recording was not proceeded. The membrane potential was maintained at around ‒80mV by injecting positive currents. Firing frequency was obtained from 1 sec step current injections (0 to +400 pA, 20 pA increment, 10 sec ISI). Bridge balance was used. The input resistance (Ri) was measured with a linear regression of voltage changes from 400 msec hyperpolarizing current injections (‒40 to ‒ 10 pA in 10 pA increments). The membrane time constant was calculated from the averaged traces to 20 consecutive hyperpolarizing current pulses (−20 pA; 400 ms) with a single exponential function (Lopez *et al*., 2012). If the series resistance (Rs) or membrane capacitance (Cm) was changed more than 20% over the recording, the cell was not further analyzed.

In voltage clamp recordings, Rs was compensated at least 60%. BK currents were measured by subtracting currents with 10 μM paxilline from the total current in the presence of 1μM tetrodotoxin (TTX) and 2 mM 4-aminopyridine (4-AP) (Montgomery and Meredith, 2012). Cells were held at −90 mV, 150 ms voltage step of ‒100 to +30 mV in 10 mV increments was applied and stepped back to ‒90 mV for 130 ms. Three current traces were averaged for analysis, and leak currents were subtracted using the P/4 method with a subsweep holding potential of −90 mV. Paxilline was applied to the slice using a local perfusion pencil for at least 10min prior to the second recording. BK current levels were obtained from the peak and normalized to cell capacitance. No paxilline-sensitive current was present in *Kcnma1*^—/—^ dentate granule neurons (n=2).

### EEG and EMG Recordings

Behavioral observations, surgeries, EEG and EMG recordings and data analyses were performed blinded to experimental condition. After daily monitoring of behavioral signs of seizures, 2-4-month-old mice were implanted with dural electrodes, with or without EMG lead implantation at the dorsal clavotrapezious neck muscles behind the base of the skull (Pinnacle Technology 4 channel EEG system, Lawrence, KS (Iffland PH, 2021). Following a minimum of 72 hours of recovery period, video-EEG recordings were obtained using Pinnacle Technology Sirenia Acquisition software for 24 hours at a sampling rate of 2 kHz. Mice were visually monitored during seizures and behavioral responses were scored using modified Racine criteria: 1) Raised tail and/or abnormal posturing; 2) Myoclonic movement of a limb, favoring one side; 3) Brief tonic-clonic convulsive episodes (approx. 1-5 sec); 4) Tonic-clonic seizures associated with rearing or jumping; and 5) Status epilepticus (Luttjohann *et al*., 2009; Van Erum *et al*., 2019). Continuous EEGs were manually reviewed for interictal epileptiform discharges and/or spontaneous seizures. Interictal epileptiform discharges were defined as discrete and sharply contoured discharges (e.g., spike and wave). Seizures were defined as at least 10 seconds of sharply contoured and rhythmic bursts of activity.

Seizure thresholds were defined in response to chemoconvulsant challenge. A baseline 15-minute video-EEG recording was obtained, followed by injection of 40 mg/kg of Pentylenetetrazol (PTZ, Sigma, Catalog# P6500, 10 mg/ml stock in sterile saline) and an additional 30 minutes recording after injection. Thirty minutes post-PTZ injection the experiment was terminated, and mice were euthanized by CO2 asphyxiation and thoracotomy. Video-EEG and EMG data were analyzed using Sirenia Seizure Pro (Pinnacle Technology, Lawrence, Kansas, USA). Experimenters were blinded to experimental group during PTZ-induced seizure, observation, and data analysis. Seizures were defined as 10 continuous seconds of sharply contoured and rhythmic discharges with a clear onset, offset, and evolution. Seizure threshold was defined as the latency to first seizure after PTZ injection. Total EEG power was reported as the sum of all frequency bands. EEG traces were examined visually for significant artifacts, and EEGs resulting in anomalous power data were excluded from the analysis, defined as EEGS with high amplitude movement artifacts (>16000µV^2^/Hz) or low amplitude signals (<750µV^2^/Hz).

EMG data were analyzed by manual review and the longest durations of attenuated EMG activity were scored. Attenuated EMG activity was defined as at least 1 second of EMG activity that was lower in amplitude than pre-PTZ injection baseline.

### Stress-induced dyskinesia assays

Two to 3-month-old mice were used in all assays for N999S and D434G cohorts. Two to 8-month-old mice were used for *Kcnma1*^—/—^ cohorts due to breeding difficulties (Meredith *et al*., 2004). For acute stress-induced dyskinesia evaluation, the total restraint time was 5 minutes.

Mice were restrained for 2.5 min by hand, clasping the dorsal cervical aspect between the index finger and thumb and the tail with the pinky finger, with the mouse dorsal side flat against the palm in a vertical upright position. Afterwards, the tail was released, leaving only the upper body restrained for 2.5 mins. Mice were then placed in a transparent 1000 ml beaker under video recording for 5 minutes. Behaviors were manually scored from videos totaled for grooming time, or the number of occurrences of rearing, sniffing, circling/hyperactivity, twisting/choreiform movement, tonic-clonic movement, flattened/dystonic posturing, tremoring, listing and falling.

Immobility/behavioral arrest was cumulatively timed from the start to cessation of each immobile episode. Behavioral parameters were modified from stereotypic behavioral scoring (Kelley, 1998) and prior dyskinesia mouse models (Khan *et al*., 2004; Khan and Jinnah, 2002; Sebastianutto *et al*., 2016; Shirley *et al*., 2008).

### Statistics

Electrophysiology and behavioral data were tested for normality with Shapiro-Wilk normality test and either parametric or non-parametric statistics were analyzed in Graphpad Prism v9.02 (San Diego, CA). Outliers were determined by the ROUT method and were included in all datasets. Data are plotted as either mean ± SEM, or individual data points with median and inter-quartile range, as indicated in figure legends. The statistical test used for each dataset is indicated in the figure legend, and *p* < 0.05 was considered significant. P-values in figure legends are reported for post hoc tests when the main effect was *p* <0.05, or reported for the main effect, if *p* >0.05. Groups with three or fewer data points were not included in statistical analysis, as noted in legends. For parametric data, two-tailed, unpaired t-tests were performed with Welch’s correction for unequal variance. For multiple comparisons, one-way ANOVAs were performed with Welch’s correction followed by Dunnett’s T3 posthoc test. Two-way repeated measures ANOVAs were performed with Geisser-Greenhouse correction followed by Bonferroni posthoc test for multiple comparisons (comparisons between genotypes across voltages). Mann-Whitney or Kruskal-Wallis followed by Dunn’s multiple comparisons were used for non-parametric data.

In gene microarray studies, differential mRNA transcript expression was determined at a 2-fold change cutoff, with *p* > 0.05 and false discovery rate, FDR = 0.99 using an ANOVA with an eBayes test was used (Ritchie *et al*., 2015)(Applied Biosystems Transcriptome Analysis Console (TAC) Software v4.0.1).

## Supporting information

Supplemental Methods, Results, Table, and Figures

Supplemental Video S2. N999S Het

Supplemental Video S3. D434G Het

Supplemental Video S5. Stress-induced dyskinesia

## ACKNOWLEDGEMENTS

This work was supported by grants from NHLBI HL102758 (A.L.M.), The Training Program in Integrative Membrane Biology NHLBI T32-GM008181 (A.L.M. and K.K.M.), The American Physiological Society’s Ryuji Ueno award sponsored by the S & R Foundation (A.L.M.), The Interdisciplinary Training Program in Muscle Biology NIAMS T32-AR007592 (S.P.), and NINDS NS114122. For gene expression studies, we acknowledge the support of the University of Maryland, Baltimore, Institute for Clinical & Translational Research (ICTR voucher #376) and the National Center for Advancing Translational Sciences (NCATS) Clinical Translational Science Award (CTSA) grant number 1UL1TR003098, Nick Ambulos and Jing Yin for performing microarray experiments, and Yuji Zhang for biostatistical analysis. We thank Huanghe Yang for discussions involving unpublished data.

## AUTHOR CONTRIBUTIONS

SMP established the transgenic lines, performed HEK cell and neuronal recordings, analyzed data, and drafted the manuscript. CER performed locomotor assays, data analysis, animal husbandry, and manuscript drafting. PHI performed EEG recordings, data analysis, and manuscript drafting. KKM assisted with EEG recordings, data analysis, and animal husbandry. HJM performed neuronal recordings, data analysis, and animal husbandry. AEP established the transgenic lines and performed animal husbandry. AGL assisted with open field assays. PBC provided essential equipment and reagents. ALM designed experiments, analyzed data, and drafted the manuscript. All authors approved the final version of the manuscript.

