## Supplemental Methods, Results, Table, and Figures for "BK channel properties correlate with neurobehavioral severity in three *KCNMA1*-linked channelopathy mouse models"

Supplemental Materials & Methods

Supplemental Results

Supplemental Figures S1-S6

Supplemental Table S1

### SUPPLEMENTAL MATERIALS AND METHODS

#### *Patch-clamp electrophysiology*

Figure S1A-C: Activation time constants were obtained from the same patches in Figure 1. $\tau_{act}$ was obtained by fitting the rising phase of the outward $K^+$ current to single exponential function. For the deactivation kinetics, BK currents were elicited by +200 mV voltage command for 20 ms from a holding potential of −100 mV followed by 15 ms voltage steps from –200 mV to –10mV with +10 mV increments. Deactivation time constants were obtained by fitting tail currents with single exponential functions. Leak currents were compensated using a P/5 protocol with a subsweep holding potential of −120 mV.

Figure S1D: BK currents were recorded in inside-out patches in physiological $K^+$ and 10 μM intracellular $Ca^{2+}$ as described in (Moldenhauer *et al.*, 2020). In voltage-clamp mode, patches were held at -150 mV, stepped from -150 mV to +150 mV for 30 ms (10 mV increments), and stepped back to -150 mV. Lisdexamfetamine dimesylate (catalog L-026, Supelco Millipore-Sigma) and dextroamphetamine (catalog 1180004, Millipore Sigma) were applied at 155 ng/ml, and paxilline (#2006; Tocris, Bristol, UK) was applied at 100 ng/ml. Current levels were assessed at baseline and 5 minutes after drug application and were normalized to control current levels for each patch.

#### *Genotyping of Kcnma1$^{N999S}$, Kcnma1$^{D434G}$, Kcnma1$^{H444Q}$ and Kcnma1$^{—/—}$ mouse lines*

Genotyping was performed tail snips by TaqMan real-time PCR at Transnetyx, Inc (Cordova, TN) using the following: *Kcnma1*$^{N999S}$ (Kcnma1-9 MUT probe set: (F) TCGGTGTTGGCTTAAGAATGCTT; (R) CCTCAGCTATTAGAGCCTCGAGCTC; WT reporter:

CAGACATACTTCAATGACAATAT; N999S reporter: CAGACATATTTCAGTGACAATAT), *Kcnma1*$^{D434G}$ (Kcnma1-8 MUT probe set: (F) CTCTAACTTCCTGAAGGACTTTCTGCACA; (R) CAGAGAGAAGCATGAGTTTAGGTGGCA; WT reporter: ACCGGGATGATGTCA; D434G reporter: ACCGTGGTGATGTCAA), and *Kcnma1*$^{H444Q}$ (Kcnma1-7 MUT probe set: (F) CTGTGGACACATTACTCTGGAGAGTG; (R) GGGTCTGGTGGCAGCA; WT reporter: TCTTACTTGTGAAGAAAG; H444Q reporter: CTCTTACTTCTGCAGAAAG).

*Kcnma1*$^{-/-}$ (Slo KO) mice were genotyped using the WT primer set (F) CATCATACCGGTGACCATGGA; (R) CCAAGAAAGCCCACCACATG; WT Reporter: CCCGGCTGTCGCACG and Neomycin primer set (F) GGGCGCCCGGTTCTT; (R) CCTCGTCCTGCAGTTCATTCA; Neo Reporter: ACCTGTCCGGTGCCC.

***Action potential waveform analysis***

Action potential (AP) amplitude was defined as the difference between the peak and threshold. Half width ($t_{1/2}$) was the width of AP at 50% of the peak amplitude from the AP threshold. The amplitude of fAHP was defined as the voltage change from the AP threshold to the most negative voltage (AHP anti-peak) after repolarization. The fAHP decay was measured as the depolarization rate from the AHP anti-peak over the first 3 ms ($\Delta V$/3ms). AP thresholds, defined as the membrane potential where the first derivative reached 10 mV/ms, were not different between genotypes for any mouse line (data not shown).

***Open Field activity***

Mice were acclimated in the testing room one hour prior to assays. Each mouse was placed in the open arena (70 x30 x25cm, (Cover *et al.*, 2019)) for 15 min. Mouse movement and total distance was analyzed in EthoVision XT (Noldus).

***Wheel Running Activity***

Mice were placed in housing cages with running wheels (Coulbourn Instruments) on a standard 12:12-hour light-dark cycle for 48 hours with *ad libitum* access to food and water. Wheel activity was measured via magnetic switches and recorded using ClockLab software (Actimetrics). Individual mouse wheel rotation counts were then quantified in 1-minute bins in ClockLab software running in Matlab v6.1 (Mathworks). The following parameters were calculated for the 12-hour dark phase as average measurements: speed, maximum speed, number of activity gaps (defined as consecutive 1-minute bins registering 0 rpm), activity gap duration, and maximum activity gap duration. All parameters were calculated by a custom python script.

***Rotarod***

Mice were acclimated to the testing environment in their normal housing cages for one hour prior to testing. Mice were trialed 3 times per day for 7 consecutive days under video capture. Body weight was measured on days 1 and 7. After placement on the rotarod (IITC Life Science Inc. Rat Mouse Rotarod), mice were acclimated on the apparatus for 30 seconds prior to 1$^{st}$ trial. During the acclimation, mice were allowed to fall off up to two times. Rod acceleration was 4 to 40 rpm over 5 minutes for each trial. Mice were given a 2-minute inter-trial interval. “Fall” was called when the mice fell off from the rod or made a 360-degree revolution around the rod. Three trials per day were averaged for each mouse.

*Hanging Wire*

Mice were acclimated to the testing environment in their home cages for one hour prior, and body weights were measured prior to the start of testing. Three consecutive trials were then performed in one day. Mice were placed right-side-up on a standard cage lid with parallel metal bars, the lid was gently shaken three times to provoke grasping the bars, then the lid was slowly inverted to put mice in the hanging position. Trial duration maximum was 120 seconds. A 10 second inter-trial interval was given after falling. Mice remaining hanging up to the maximum time were inverted to right-side-up on the lid and given a 10 second interval before the next trial.

### SUPPLEMENTAL RESULTS

#### Effects of GOF (N999S and D434G) and LOF (H444Q) variants on BK current kinetics

In addition to shifting the voltage dependence of activation, variants were also evaluated for their effects BK channel opening and closing rates. Activation kinetics were obtained from fits of the rising phase of the macroscopic current at each voltage step when channels are opening before they reach steady-state ($\tau_{act}$, Figure S1B). $BK^{WT}$ current activation ranged from 8.2 ± 1.12 to 9.5 ± 1.4 ms at 100 to 130 mV, decreasing with more positive voltage steps to 1.2 ± 0.1 ms at 250 mV. $BK^{N999S}$ and $BK^{D434G}$ currents had faster activation compared to $BK^{WT}$, at all voltages ($BK^{N999S}$) or above 120 mV ($BK^{D434G}$). At lower voltages, $BK^{D434G}$ channels were more steeply voltage dependent, but did not exceed the fast activation time constants of $BK^{N999S}$ channels. At +170 mV (Figure S1B inset), $\tau_{act}$ for $BK^{WT}$ currents was 3.8 ± 0.3 ms, while $BK^{N999S}$ and $BK^{D434}$ currents activated faster (0.9 ± 9.1 ms and 1.8 ± 0.1 ms, respectively). $BK^{H444Q}$ current activation did not differ from $BK^{WT}$ until voltages above 160 mV, which is near

$V_{1/2}$ of $BK^{H444Q}$. At 160 mV, $\tau_{act}$ for $BK^{H444Q}$ (7 ± 1 ms) was slower than $BK^{WT}$ (5 ± 0.4 ms). At 170 mV (Figure S1B inset), $\tau_{act}$ for $BK^{H444Q}$ (6.5 ± 0.5 ms) was slower than $BK^{WT}$ (3.8 ± 0.3 ms). Therefore, GOF effects were introduced into activation kinetics by the N999S and D434G variants, while H444Q conferred LOF effects.

To measure closing rates, channels were first opened with a depolarizing step to 200 mV, and then closed with steps to varying voltages (–200 to –50 mV). Deactivation kinetics were obtained from fits of the falling phase of the macroscopic tail current ($\tau_{deact}$, Figure S1C). $BK^{WT}$ deactivation kinetics showed a small increase at more positive voltages, from 0.1 ± 0.01 ms at –200 mV to 0.7 ± 0.01 ms at –20 mV. Variant-containing channels followed the same trend with slower closing at more depolarized voltages. However, $BK^{N999S}$ and $BK^{D434}$ currents deactivated slower than $BK^{WT}$ across the voltage range (4.5 ± 0.7 ms and 1.5 ± 0.1 ms at –20 mV, respectively). $BK^{H444Q}$ deactivated more quickly than $BK^{WT}$ between –140 mV to –20 mV (0.4 ± 0.01 ms at –20 mV). Slower channel closure is consistent with GOF effects (N999S and D434G), while faster closing denotes LOF (H444Q).

**Effects of dextroamphetamine and lisdexamfetamine on $BK^{WT}$ and $BK^{N999S}$ channels**

Dextroamphetamine (D-amphetamine) and lisdexamfetamine, a pro-drug of D-amphetamine and L-lysine, have been reported to treat PNKD episodes in six children harboring N999S variants, as well as one child harboring another GOF variant, N536H (Keros *et al.*, 2021; Zhang *et al.*, 2020). A related stimulant, methamphetamine was proposed to inhibit BK channels through multiple mechanisms in dopaminergic neurons (Lin *et al.*, 2016; Tatro *et al.*, 2013; Wang *et al.*, 2013), raising the question of whether dextroamphetamine and lisdexamfetamine can modulate BK channel activity. To test if either drug has a direct effect on BK currents in heterologous cells, lisdexamfetamine and dextroamphetamine were applied to patches from HEK293 cells expressing $BK^{WT}$ or $BK^{N999S}$ (Figure S1D). Macroscopic BK currents were evaluated after perfusion of each drug and compared to pre-drug control current levels. No inhibition of current level was observed for either $BK^{WT}$ or $BK^{N999S}$ patches with either drug. However, the inhibitor Paxilline fully abrogated $BK^{WT}$ or $BK^{N999S}$ currents at the end of each experiment.

**Passive membrane properties and action potential waveforms**

Resting membrane potential (RMP) and input resistance ($R_i$) parameters were first assessed to determine whether changes in passive membrane properties contributed to increased firing. From the same dentate granule neurons used for action potential recording, RMP was measured in current-clamp mode prior to any current injection, and $R_i$ was measured from the voltage offset in response to hyperpolarizing current injections (Table S1). RMPs across WT controls were –80 to –82 mV. No significant depolarizations were observed in any of the transgenic neurons. Similarly, the range for $R_i$ (320 to 364 MΩ for WT neurons) was not different in any transgenic condition. These data are consistent with the lack of BK current activation in this range in voltage-clamp recordings for $Kcnma1^{N999S/WT}$, $Kcnma1^{D434G/WT}$ or $Kcnma1^{H444Q/WT}$ (Figure 2) and argue against changes in the subthreshold membrane sensitivity to current injection initiating the increased firing with the GOF variants. Previous studies in dentate granule cells found no effect of BK channel inhibition on RMP or $R_i$ (Bock and Stuart, 2016; Brenner *et al.*, 2005). In contrast, increased BK channel activation in two other GOF mouse lines, $Tg\text{-}BK^{R207Q}$ and $\beta2^{-/-}$, both hyperpolarized the baseline membrane potentials of spontaneously firing neurons and decreased $R_i$ (Montgomery *et al.*, 2013; Whitt *et al.*, 2016), though these changes oppose the direction expected for increased excitability.

The only condition showing a difference from the respective WT control was a greater membrane capacitance ($C_m$) in *Kcnma1*$^{D434G/WT}$ neurons (Supplemental Table 1). This difference could have the potential to reduce the firing gain at lower current injections, preventing the *Kcnma1*$^{D434G/WT}$ I-O curve from looking similar to *Kcnma1*$^{N999S/WT}$. However, an explanation for this change in $C_m$ is unclear from the data.

To examine whether the action potential revealed any differences between *Kcnma1*$^{N999S/WT}$ and *Kcnma1*$^{D434G/WT}$ neurons, waveforms were analyzed from experiments in Figure 3. Modulation of action potential repolarization and AHP in repetitive firing occurs with both BK channel inhibition (slower repolarization and reduced AHP amplitude) and activation (faster repolarization and increased AHP amplitude) (Dong *et al.*, 2021; Gu *et al.*, 2007; Montgomery and Meredith, 2012; Shao *et al.*, 1999). Neurons from a separately generated *Kcnma1*$^{D434G}$ mouse line and from SLO-E366G flies showed reduced action potential width and increased AHP amplitude (Dong *et al.*, 2021; Kratschmer *et al.*, 2021), predicting that the firing increases resulting from N999S and D434G would be a consequence of decreased action potential half-width ($t_{1/2}$) and/or increased fAHP amplitude.

To test this, action potential waveforms were analyzed at the 200 pA current injection step, where firing rate was increased for *Kcnma1*$^{N999S/WT}$ and *Kcnma1*$^{D434G/D434G}$ but not for *Kcnma1*$^{D434G/WT}$ (Figure S3). In *Kcnma1*$^{WT/WT}$ neurons, the action potential half-width was 2.4 ± 0.1 ms, and the fAHP amplitude was 12 ± 0.7 mV (Figure S3A-C). *Kcnma1*$^{N999S/WT}$ action potentials were indistinguishable in these parameters. This result was unexpected, given the large hyperpolarizing shift in the $V_{1/2}$ and faster activation observed from BK$^{N999S}$ channels compared to WT (Figure 1C-D, S1B). Instead, the AHP decay was steeper than WT controls (*Kcnma1*$^{N999S/WT}$ 0.60±0.03 mV/ms compared to *Kcnma1*$^{WT/WT}$ 0.41±0.03 mV/ms; Figure S3D). The faster depolarization rate would facilitate more rapid subsequent action potential initiation. However, because BK$^{N999S}$ channels actually exhibit slower deactivation than WT (Figure S1C), which would be expected to prolong the AHP, the effect may be indirect.

Like the changes in *Kcnma1*$^{N999S/WT}$, *Kcnma1*$^{D434G/D434G}$ waveforms were altered in a similar manner in comparison to their respective control neurons. No changes were found in action potential half-width or AHP amplitude (Figure S3E-G), but the AHP decay rate was faster (0.69±0.04 mV/ms and 0.46±0.04 mV/ms, respectively; Figure S3H) suggesting a similar mechanistic basis for the increased *Kcnma1*$^{D434G/D434G}$ firing rate. BK$^{D434G}$ channels also activated faster, with a more hyperpolarized $V_{1/2}$, and deactivated more slowly than WT (Figure 1C, S1B-C), pointing to a discrepancy in the direct translation of channel properties recorded in heterologous cells into the dynamic neuronal activity context.

Since *Kcnma1*$^{D434G/WT}$ neurons do not show increased firing at 200 pA, these action potentials would not be expected to differ in parameters related to setting firing frequency. Consistent with the hypothesis that AHP decay rate sets the firing frequency, *Kcnma1*$^{D434G/WT}$ values were similar to WT (0.56±0.02 mV/ms, Figure S3H). However, *Kcnma1*$^{D434G/WT}$ neurons did have shorter half-widths, an effect that was not observed in either *Kcnma1*$^{N999S/WT}$ or *Kcnma1*$^{D434G/D434G}$ neurons. Altogether, these results demonstrate that BK channels regulate multiple phases of the action potential in dentate granule neurons, and further suggest that AHP decay is a more important determinant of frequency than repolarization under these firing conditions.

**Additional dyskinesia and motor impairment caused by N999S and D434G**

Voluntary wheel running has a positive emotive valence for mice and was used as a secondary investigation for PNKD-like behavior. Mice produced stereotypical nocturnal wheel activity patterns during the 48-hr assay. Episodic immobility or hypotonia would be predicted to reduce running. *Kcnma1*$^{N999S/WT}$ mice showed decreased average distance during the assessment period (Supplemental Figure S5Ai). The reduction in running distance was not due to a decrease in the maximum speed the mice could achieve, but instead occurred as a result of increased gaps in activity (Supplemental Figure S5Bi, Ci). These gaps could be consistent with cessation of running during a PNKD-like immobility, similar to that observed in the beaker after stress. However, we cannot rule out reduced distance as a result of a baseline dyskinesia or muscle weakness.

Also similar to the beaker test, *Kcnma1*$^{D434G/WT}$ (*Kcnma1*$^{D434G/D434G}$ were not available for this assay), *Kcnma1*$^{H444Q/WT}$, and *Kcnma1*$^{H444Q/H444Q}$ mice did not show any difference in distance, although *Kcnma1*$^{D434G/WT}$ mice did have lower maximum speeds and longer gap durations (Supplemental Figure S5Bii-iii, Cii-iii, Dii-iii). Whether the changes observed in *Kcnma1*$^{D434G/WT}$ running patterns are also consistent with either a baseline dyskinesia or a PNKD-like episode cannot be unambiguously delineated from this data. Although running wheel performance is dependent on more factors than the motor function displayed in the beaker, it provides a second line observation consistent with the interpretation of PNKD caused by N999S in response to both negative (stress) and positive (running) triggers.

*Kcnma1*$^{-/-}$ mice were also included in the voluntary wheel running assay as a control because they have been well-characterized to exhibit baseline ataxia, gait impairment, and reduced strength and wheel running (Meredith *et al.*, 2004; Meredith *et al.*, 2006; Sausbier *et al.*,

2004). Consistent with previous data, *Kcnma1*$^{-/-}$ mice showed reduced distances covered, reduced maximum speed, and longer activity gap durations compared to WT littermates (Supplemental Figure S5A-Div). This phenotype was distinct from the changes observed in *Kcnma1*$^{N999S/WT}$ mice, suggesting that the latter's reduction in distance may not be related to hypokinetic activity, as in the case of ataxic mice like *Kcnma1*$^{-/-}$. *Kcnma1*$^{-/+}$ mice did not show any significant differences from WT littermates.

Mice were further evaluated for baseline dyskinesia and motor coordination abnormalities in additional behavioral assays. Among patients, multiple motor abnormalities have been described in addition to PNKD including dystonia/chorea and myoclonus, hypotonia and weakness, dyspraxia and ataxia, motor reflex impairment, and tremor (Heim *et al.*, 2020; Miller *et al.*, 2021; Wang *et al.*, 2017; Zhang *et al.*, 2015). BK channels are prominently expressed in key motor regulatory areas such as somatosensory cortex, basal ganglia, and cerebellum (Bailey *et al.*, 2019; Sausbier *et al.*, 2005). In cerebellar function, both *Kcnma1*$^{-/-}$ and another *Kcnma1*$^{D434G}$ mouse lines show impaired behavioral motor performance, with decreased and increased firing in cerebellar Purkinje cells, respectively (Dong *et al.*, 2021; Sausbier *et al.*, 2004). *Kcnma1*$^{-/-}$ mice also have altered neuromuscular junction function associated with muscle weakness (Wang *et al.*, 2020). Baseline changes in motor coordination and/or grip strength are relatively common in other paroxysmal dyskinesia mouse models as well (Khan *et al.*, 2004; Khan and Jinnah, 2002; Pan *et al.*, 2020; Tan *et al.*, 2018). To further understand the background for the dyskinesia more completely, additional locomotor assays were performed to assess the degree of baseline motor impairment.

Acute muscle strength was first tested by hanging mice from a stationary platform (cage lid) for 120 sec. Most WT mice can hang on for several minutes before losing grip and falling

(Crawley, 2000), although each WT control cohort exhibited a wide range of latency to fall values. *Kcnma1*$^{N999S/WT}$ mice fell off earlier than WT littermates indicating weaker grip strength (Figure S6A). However, *Kcnma1*$^{D434G/WT}$ and *Kcnma1*$^{D434G/D434G}$ mice had no differences in fall latencies, providing no evidence for acute differences in strength (Figure S6B). *Kcnma1*$^{H444Q/H444Q}$ mice showed shorter fall latencies than *Kcnma1*$^{H444Q/WT}$ mice, which were used as controls due to the shortage of WT littermates (Figure S6C). *Kcnma1*$^{-/-}$ mice had the shortest fall latencies (Figure S6D), consistent with previous reports (Meredith *et al.*, 2004; Sausbier *et al.*, 2004; Wang *et al.*, 2020; Yao *et al.*, 2021). No *Kcnma1*$^{-/-}$ mouse had the ability to hang on longer than 60 sec, which was a tighter cluster than the spread in *Kcnma1*$^{N999S/WT}$ values. Because of this, it seems unlikely that the hanging wire assay was a significant trigger for PNKD-like immobility in *Kcnma1*$^{N999S/WT}$ mice, potentially due to the short duration compared to the full restraint stress used in Figure 7. The results give the series of *Kcnma1*$^{-/-}$ > *Kcnma1*$^{N999S/WT}$ > *Kcnma1*$^{H444Q/H444Q}$ > *Kcnma1*$^{D434G/D434G}$ for muscle weakness.

Motor coordination was tested by rotarod over 7 days of trials on an accelerating rod. Hypotonia and PNKD-like paroxysms would be expected to produce extremely short latencies to fall, since a single major loss of tone would be catastrophic for coordination. Alternately, baseline coordination could be impaired in the absence of immobility. Over the week of trials, *Kcnma1*$^{N999S/WT}$ mice showed shorter fall latency compared to their WT littermates, with the decrease on the last day, but not the first, being statistically significant (Figure S6E). Motor learning was observable as an improvement in fall latency times across the 3 trials on each day (data not shown), suggesting the overall impairment was related to motor coordination and not learning. Among the individual mice, there was no explicit evidence for individual trials with extremely short fall latencies that would be consistent with the triggering of sudden hypotonic events. However, partial loss of tone might be compensated for by the high level of attention induced in this assay. Compared to the voluntary running wheel activity, successful navigation of the rod's surface and rotation requires a higher degree of motor coordination than the wider home-cage wheel. Mice fall off the rotarod at lower speeds than the maximum speeds achieved on the voluntary running wheel.

In contrast, *Kcnma1*$^{D434G/WT}$ mice did not show any difference in fall latency or motor learning compared to their WT controls (Figure 8F). However, homozygous *Kcnma1*$^{D434G/D434G}$ had large reductions in both. *Kcnma1*$^{H444Q/WT}$ and *Kcnma1*$^{H444Q/H444Q}$ performance was highly variable, reducing the ability to make a firm conclusion from this data (Figure S6G). *Kcnma1*$^{—/—}$ mice performed the worst, with extremely short latencies to fall indicative of their profound ataxia (Figure S6H). The motor coordination severity fell in the series *Kcnma1*$^{—/—}$ > *Kcnma1*$^{D434G/D434G}$ > *Kcnma1*$^{N999S/WT}$ > *Kcnma1*$^{H444Q/H444Q}$ for motor coordination and learning.

**SUPPLEMENTAL ACKNOWLEDGEMENTS**

We thank Todd Gould, Junfang Wu, and Brian N. Mathur for generously providing use of equipment for mouse motor assays and helpful discussions.

### Supplemental Figure Legends

#### Table S1. Passive membrane properties

Resting membrane potential, input resistance, membrane time constant, and capacitance from *Kcnma1*$^{N999S/WT}$, *Kcnma1*$^{D434G/D434G}$, *Kcnma1*$^{H444Q/WT}$, and *Kcnma1*$^{H444Q/H444Q}$ dentate granule neurons were not significantly different from their respective controls (P > 0.05, unpaired t-test and One-way ANOVA, respectively). *Capacitance was greater from *Kcnma1*$^{D434G/WT}$ neurons compared to WT ($p$=0.0036). Data are presented as mean ± SEM.

#### Figure S1. BK current properties recorded in HEK cells

(A) Representative macroscopic BK currents recorded from inside–out patches in symmetrical $K^+$ and 1 μM intracellular $Ca^{2+}$; analysis from data recorded in main Figure 1C-D. Patches were held at −100 mV, stepped from −100 mV to 250 mV for 30 ms, followed by a tail step −100 mV for 15 ms. Scale bars: 1 nA, 5 ms. (B) Activation time constants ($\tau_{act}$), obtained from single exponential fits of the current rising phase from 100 mV to 250 mV. BK$^{N999S}$ and BK$^{D434G}$ channels had decreased $\tau_{act}$ compared to BK$^{WT}$, either across all voltage steps (Mixed effects model for repeated measures with Bonferroni posthoc, $p$<0.01) or between 120 mV to 250 mV ($p$<0.05), respectively. BK$^{H444Q}$ channels had had increased $\tau_{act}$ compared to BK$^{WT}$ between 160 mV to 250 mV ($p$<0.05). *Inset*: Representative current traces from 170 mV step, scaled to the maximal current to illustrate activation timecourse. Scale bar: 10 ms. (C) Deactivation time constants ($\tau_{deact}$), obtained after an activating step to 200 mV (20 ms) from single exponential fits of the tail step current decay elicited from –200 to –50 mV (for 15 ms). BK$^{N999S}$ and BK$^{D434G}$ channels had increased $\tau_{deact}$ compared to BK$^{WT}$, across all voltage steps (Mixed effects model for repeated measures with Bonferroni posthoc, $p < 0.01$), with the exception of –160 ($p > 0.05$), respectively. $BK^{H444Q}$ channels had had decreased $\tau_{deact}$ compared to $BK^{WT}$ between –190 mV and between –140 and –20 mV ($p < 0.05$). *Inset*: Representative current traces from –40 mV step, scaled to the maximal current to illustrate deactivation timecourse. Scale bar: 0.5 ms. Data are presented as mean ± SEM. (D) Effects of 155ng/mL lisdexamfetamine (lis) and dextroamphetamine (d-amp) on $BK^{WT}$ and $BK^{N999S}$ channels. Normalized data presented as the proportion of the maximal current ($I/I_{max}$) for each patch, before (Control) and after drug application. Neither lis ($BK^{N999S}$: n = 7, P= 0.98; One-way ANOVA) nor d-amp ($BK^{WT}$: n = 4, P= 0.99; One-way ANOVA and $BK^{N999S}$: n = 6, P= 0.15; One-way ANOVA) produced a decrease in BK current levels, while 100 nM paxilline (pax) inhibited both $BK^{WT}$ and $BK^{N999S}$ currents (P< 0.001; One-way ANOVA for all).

**Figure S2. Introduction of patient variants into the mouse *Kcnma1* gene by CRISPR/Cas9 editing.** (A) *Kcnma1*$^{N999S/WT}$ mice were generated by introducing a non–synonymous mutation within the codon AAT→ A**g**T (red boxes) in exon 25. WT sequence is C57BL/6J. Underlined nucleotides are the gRNA sequence. Lowercase letters denote mutations. Chromatogram from an N1 heterozygous mouse. (B) *Kcnma1*$^{D434G/WT}$ mice were generated by mutation within the codon GAT→ G**g**T in exon 10. Chromatogram from a founder mouse. (C) *Kcnma1*$^{H444Q/WT}$ mice were generated by mutation within the codon CAC → CA**g** in exon 10 (same guide RNA as D434G). Chromatogram from a founder mouse.

**Figure S3. *Kcnma1*$^{N999S/WT}$, *Kcnma1*$^{D434G/WT}$, and *Kcnma1*$^{D434G/D434G}$ action potential waveforms**

(A) Superimposed *Kcnma1*$^{N999S/WT}$ and *Kcnma1*$^{WT/WT}$ waveforms from the 10$^{th}$ action potential at the 200pA current injection from data in Figure 4. (B) Action potential half-width ($t_{1/2}$) was not different in *Kcnma1*$^{N999S/WT}$ neurons versus WT controls (*p*=0.6617; Mann-Whitney test). (C) fAHP amplitude was not different in *Kcnma1*$^{N999S/WT}$ neurons versus WT controls (*p*=9214; Mann-Whitney test). (D) AHP decay 3 ms after the peak was faster in *Kcnma1*$^{N999S/WT}$ neurons compared to WT controls (**p*=0.0002; t-test). (E) Superimposed *Kcnma1*$^{D434G/D434G}$, *Kcnma1*$^{D434G/WT}$ and *Kcnma1*$^{WT/WT}$ waveforms from the 10$^{th}$ action potential at the 200pA current injection from data in Figure 4. (F) $t_{1/2}$ was different in *Kcnma1*$^{D434G/WT}$ (**p*=0.0002; Kruskal-Wallis test), but not *Kcnma1*$^{D434G/D434G}$ neurons (*p*=0 >0.9999). (G) fAHP amplitudes were comparable in *Kcnma1*$^{D434G}$ mice (*p*=0.3441; Kruskal-Wallis test). (H) AHP decay 3 ms after the peak was faster in *Kcnma1*$^{D434G/D434G}$ neurons compared to WT controls (**p*=0.0002), but not in *Kcnma1*$^{D434G/WT}$(*p*=0.0620; One-way ANOVA). Data are presented as mean ± SEM, with individual data points.

**Figure S4. Open field.** *Kcnma1*$^{N999S/WT}$ mice (n=8) covered the same distance as *Kcnma1*$^{WT/WT}$ mice (n=8) in a 15-minute trial (*p*=0.6973; t-test). Data are presented as individual data points with median and inter-quartile range.

**Figure S5. Locomotor wheel running. Parameters calculated from average activity counts over 48 hrs from singly housed mice with free access to wheels.** Data are presented as individual data points with median and inter-quartile range. (A*i*) Distance covered was reduced for *Kcnma1*$^{N999S/WT}$ (n=12) compared to *Kcnma1*$^{WT/WT}$ mice (n=11, **p*=0.0411; t-test). (A*ii*) Distance was comparable between *Kcnma1*$^{D434G/WT}$ (n=11) and *Kcnma1*$^{WT/WT}$ mice (n=12, *p*=0.8118; t-test). (A*iii*) Distance was comparable between *Kcnma1*$^{H444Q/H444Q}$ (n=7) and *Kcnma1*$^{H444Q/WT}$ (n=6, *p*=0.4880; t-test). *Kcnma1*$^{WT/WT}$ mice were not included in statistical analysis in panels A*iii*-D*iii* due to small sample size. (A*iv*) Distance was reduced for *Kcnma1*$^{-/-}$ (n=10) compared *Kcnma1*$^{+/+}$ (n=9, **p*=0.0032), but not for *Kcnma1*$^{-/+}$ mice (n=15, *p*=0.9057; One-way ANOVA). (B*i*) Maximum speed was comparable between *Kcnma1*$^{N999S/WT}$ (n=11) and *Kcnma1*$^{WT/WT}$ mice (n=12, , *p*=0.3618; t-test). (B*ii*) Maximum speed was lower for *Kcnma1*$^{D434G/WT}$ (n=11) compared to *Kcnma1*$^{WT/WT}$ mice (n=12, **p*=0.0085; t-test). (B*iii*) Maximum speed was comparable between *Kcnma1*$^{H444Q/H444Q}$ (n=7) and *Kcnma1*$^{H444Q/WT}$ (n=6, *p*=0.3634; t-test). (B*iv*) Maximum speed was lower for *Kcnma1*$^{-/-}$ (n=10) compared to *Kcnma1*$^{+/+}$ (n=9, **p*=0.0024), but not for *Kcnma1*$^{-/+}$ mice (n=15, *p*=0.9871; One-way ANOVA). (C*i*) Duration of time off wheels (gap duration) was comparable between *Kcnma1*$^{N999S/WT}$ (n=11) and *Kcnma1*$^{WT/WT}$ mice (n=12, *p*=0.8281; t-test). (C*ii*) Gap duration was reduced for *Kcnma1*$^{D434G/WT}$ (n=11) compared to *Kcnma1*$^{WT/WT}$ mice (n=12, **p*=0.0467; t-test). (C*iii*) Gap duration was comparable between *Kcnma1*$^{H444Q/H444Q}$ (n=7) and *Kcnma1*$^{H444Q/WT}$ mice (n=6, *p*=0.8326; t-test). (C*iv*) Gap duration was higher in *Kcnma1*$^{-/-}$ (n=10) compared to *Kcnma1*$^{+/+}$ mice (n=9, **p*=0.0026), but not for *Kcnma1*$^{-/+}$ mice (n=15, *p*=0.8987; One-way ANOVA). (D*i*) Number of times the mouse was off the wheel (gap events) was higher for *Kcnma1*$^{N999S/WT}$ (n=12) compared to *Kcnma1*$^{WT/WT}$ mice (n=11, **p*=0.0040; t-test). (D*ii*) Gap events were comparable between *Kcnma1*$^{D434G/WT}$ (n=11) and *Kcnma1*$^{WT/WT}$ mice (n=12, *p*=0.7425; t-test). (D*iii*) Gap events were comparable between *Kcnma1*$^{H444Q/H444Q}$ (n=7) and *Kcnma1*$^{H444Q/WT}$ (n=6, *p*=0.9341; t-test). (D*iv*) Gap events were comparable for *Kcnma1*$^{-/-}$ (n=10), *Kcnma1*$^{-/+}$ (n=15), and *Kcnma1*$^{+/+}$ mice (n=9, *p*=0.3047; One-way ANOVA).

**Figure S6. Hanging wire and rotarod.** (A-D) Time to fall in the hanging wire assay. Data are presented as individual data points with median and inter-quartile range. (A) Fall latency was lower in *Kcnma1*$^{N999S/WT}$ (n=11) compared to *Kcnma1*$^{WT/WT}$ mice (n=10, **p*=0.0014; Mann-Whitney test). (B) Fall latency was comparable for *Kcnma1*$^{D434G/D434G}$ (n=4), *Kcnma1*$^{D434G/WT}$ (n=11), and *Kcnma1*$^{WT/WT}$ mice (n=11, *p*=0.8329; Kruskal-Wallis test). (C) Fall latency was reduced for *Kcnma1*$^{H444Q/H444Q}$ (n=8) compared to *Kcnma1*$^{H444Q/WT}$ mice (n=14, **p*=0.0465; Mann-Whitney test). *Kcnma1*$^{WT/WT}$ mice were not included in the statistical analysis due to small sample size (n=3). (D) Fall latency times were lower for *Kcnma1*$^{-/-}$ mice (n=10) compared to *Kcnma1*$^{+/+}$ mice (n=10, **p*=0.0036), but not for *Kcnma1*$^{-/+}$ mice (n=17, *p*>0.9999; Kruskal-Wallis test). (E-H) Time to fall in rotarod assay. Data are presented as mean ± SEM. *, $p < 0.05$, repeated measures ANOVA with Bonferroni posthoc. (E) Fall latency was lower for *Kcnma1*$^{N999S/WT}$ mice (n=11) on day 2 (*p*=0.0045) and day 7 (*p*=0.0124) compared to *Kcnma1*$^{N999S/WT}$ mice (n=12) (indicated with *). (F) Fall latency was lower for *Kcnma1*$^{D434G/D434G}$ mice (n=4) on day 2 (*p*<0.0001), day 4 (*p*=0.0030), day 6 (*p*<0.0001) and day 7 (*p*=0.0009) compared to *Kcnma1*$^{WT/WT}$ mice (n=21) (*), but were comparable in *Kcnma1*$^{D434G/WT}$ mice (n=18). (G) Fall latency was comparable between *Kcnma1*$^{H444Q/H444Q}$ (n=7) and *Kcnma1*$^{H444Q/WT}$ (n=10). *Kcnma1*$^{WT/WT}$ mice were not included in the statistical analysis due to small sample size (n=2). (H) Fall latency was lower for *Kcnma1*$^{-/-}$ mice (n=6) on day 1 (*p*=0.0277), day 2 (*p*=0.0056), day 3 (*p*=0.0122), day 4 (*p*=0.0081), day 5 (*p*=0.0166), day 6 (*p*=0.0071) and day 7 (*p*=0.0168) compared to *Kcnma1*$^{+/+}$ mice (n=6) (*), but were comparable in *Kcnma1*$^{-/+}$ mice (n=13).

**Supplemental Videos S1-S4.** Video-EEG segments of initial seizures approximately 3 min after PTZ injection played at 1X speed.

**S1. *Kcnma1*$^{WT/WT}$.** The mouse displays clonic extensions of its hindlimbs accompanied by brief generalized epileptiform discharges on EEG (< 2 sec bursts) (t = ~20 sec). As the video progresses, epileptiform bursts increase in frequency and duration, but the mouse does not develop tonic-clonic seizures. This video represents a typical response for WT animals and illustrates the baseline behavioral changes to a 40 mg/kg PTZ dose.

**S2. *Kcnma1*$^{N999S/WT}$.** This mouse begins having burst of epileptiform discharges on EEG that are accompanied by clonic extension of the hindlimbs and brief myoclonic jerking movements (t = ~10 sec). Discharges become more sharply contoured, increase in frequency, and are generalized at t = ~30 sec. Following this (t = ~53 sec), the discharges evolve into a generalized tonic-clonic seizure lasting 13 seconds. After the seizure, the EEG is suppressed with intermittent epileptiform bursts lasting approximately 1 second.

**S3. *Kcnma1*$^{D434G/WT}$.** The video starts with the mouse in behavioral arrest that progresses to brief tonic-clonic seizures (t = ~30 sec). Tonic-clonic seizures increase in duration and continue until the end of the video segment. Between seizures, there is severe EEG suppression (t = 55 sec).
S4. *Kcnma1*$^{—/—}$. The video starts with the mouse having bursts of epileptiform discharges without clear behavioral correlates. At t = ~20 sec, the mouse has an electrographic seizure with minimal behavioral changes that include myoclonic extension of the hind limbs and occasional myoclonic jerks. Thereafter, discharges continue on EEG but are accompanied by significant EMG suppression (bottom, red). By t= 58 sec, electrographic seizures are longer in duration, and bursts of activity are higher in frequency. However, the EMG remains suppressed and the animal is shown in behavioral arrest with myoclonic extension of the hind limbs. As the seizure progresses (t = 90 sec), the mouse walks in the cage, but no abnormal tonic-clonic activity is observed.

**Supplemental Videos S5. Restraint stress-induced dyskinesia.** Five minutes of beaker activity for *Kcnma1*$^{N999S/WT}$, *Kcnma1*$^{WT/WT}$, and *Kcnma1*$^{-/-}$ mice after 5 min restraint stress. Video is a representative one-minute segment for each mouse played at 1X speed.

| | N | Resting membrane potential (mV) | Input resistance (MΩ) | Membrane time constant (ms) | Membrane capacitance (pF) |
|---|---|---|---|---|---|
| *Kcnma1*$^{WT/WT}$ | 16 | -80 ± 1 | 324 ± 12 | 24 ± 1 | 76 ± 2 |
| *Kcnma1*$^{N999S/WT}$ | 23 | -80 ± 1 | 351 ± 13 | 25 ± 1 | 72 ± 2 |
| *Kcnma1*$^{WT/WT}$ | 22 | -82 ± 1 | 364 ± 15 | 27 ± 1 | 75 ± 2 |
| *Kcnma1*$^{D434G/WT}$ | 27 | -83 ± 1 | 344 ± 11 | 30 ± 1 | 86 ± 2* |
| *Kcnma1*$^{D434G/D434G}$ | 19 | -82 ± 1 | 371 ± 14 | 27 ± 1 | 74 ± 3 |
| *Kcnma1*$^{WT/WT}$ | 7 | -80 ± 2 | 320 ± 18 | 22 ± 2 | 68 ± 3 |
| *Kcnma1*$^{H444Q/WT}$ | 8 | -82 ± 1 | 342 ± 35 | 22 ± 2 | 67 ± 5 |
| *Kcnma1*$^{H444Q/H444Q}$ | 11 | -79 ± 1 | 345 ± 16 | 22 ± 1 | 65 ± 3 |

Fig S1

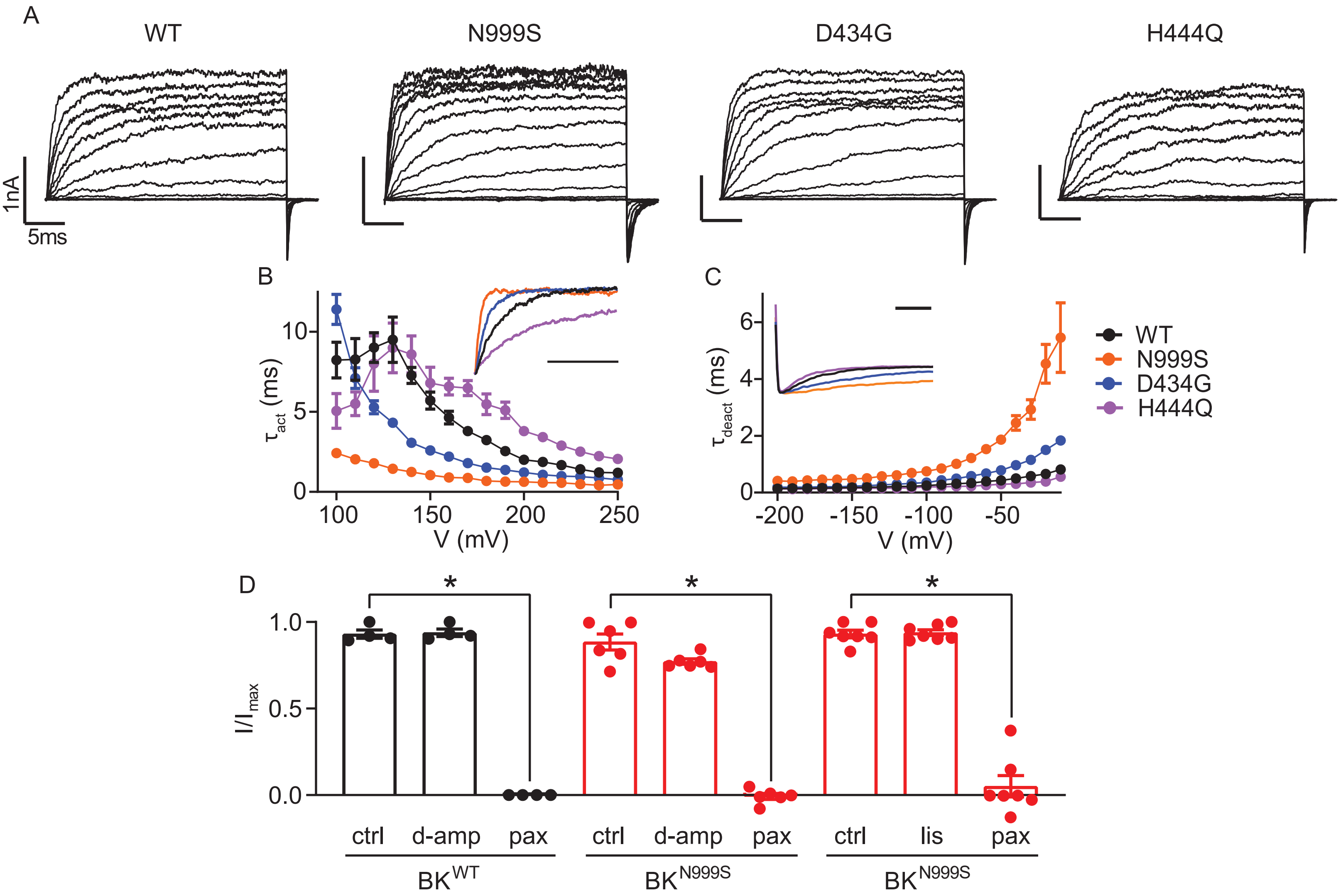

A
WT
N999S
D434G
H444Q
1nA
5ms
B
$\tau_{act}$ (ms)
10
5
0
100
150
200
250
V (mV)
C
$\tau_{deact}$ (ms)
6
4
2
0
-200
-150
-100
-50
V (mV)
WT
N999S
D434G
H444Q
D
$I/I_{max}$
1.0
0.5
0.0
*
*
*
ctrl
d-amp
pax
ctrl
d-amp
pax
ctrl
lis
pax
$BK^{WT}$
$BK^{N999S}$
$BK^{N999S}$

Fig S2

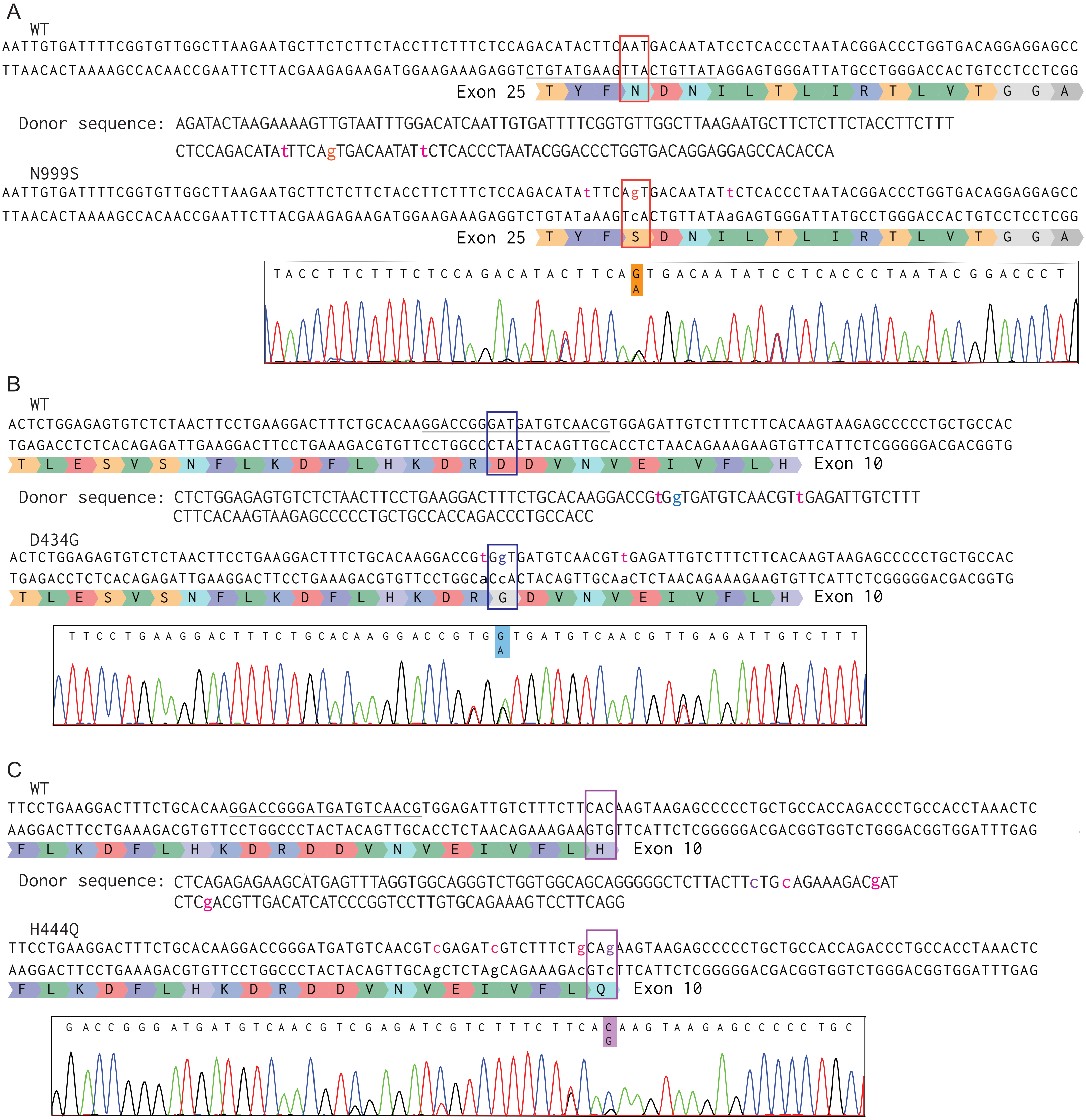
A
WT
AATTGTGATTTTCGGTGTTGGCTTAAGAATGCTTCTCTTCTACCTTCTTTCTCCAGACATACTTCAATGACAATATCCTCACCCTAATACGGACCCTGGTGACAGGAGGAGCC
TTAACACTAAAAGCCACAACCGAATTCTTACGAAGAGAAGATGGAAGAAAGAGGTCTGTATGAAGTTACTGTTATAGGAGTGGGATTATGCCTGGGACCACTGTCCTCCTCGG
Exon 25 T Y F N D N I L T L I R T L V T G G A
Donor sequence: AGATACTAAGAAAAGTTGTAATTTGGACATCAATTGTGATTTTCGGTGTTGGCTTAAGAATGCTTCTCTTCTACCTTCTTT
CTCCAGACATAtTTCAgTGACAATATtCTCACCCTAATACGGACCCTGGTGACAGGAGGAGCCACACCA
N999S
AATTGTGATTTTCGGTGTTGGCTTAAGAATGCTTCTCTTCTACCTTCTTTCTCCAGACATAtTTCAgTGACAATATtCTCACCCTAATACGGACCCTGGTGACAGGAGGAGCC
TTAACACTAAAAGCCACAACCGAATTCTTACGAAGAGAAGATGGAAGAAAGAGGTCTGTATaAAGTcACTGTTATAaGAGTGGGATTATGCCTGGGACCACTGTCCTCCTCGG
Exon 25 T Y F S D N I L T L I R T L V T G G A
TACCTTCTTTCTCCAGACATACTTCAGTGACAATATCCTCACCCTAATACGGACCCT
A
B
WT
ACTCTGGAGAGTGTCTCTAACTTCCTGAAGGACTTTCTGCACAAGGACCGGGATGATGTCAACGTGGAGATTGTCTTTCTTCACAAGTAAGAGCCCCCTGCTGCCAC
TGAGACCTCTCACAGAGATTGAAGGACTTCCTGAAAGACGTGTTCCTGGCCCTACTACAGTTGCACCTCTAACAGAAAGAAGTGTTCATTCTCGGGGGACGACGGTG
T L E S V S N F L K D F L H K D R D D V N V E I V F L H Exon 10
Donor sequence: CTCTGGAGAGTGTCTCTAACTTCCTGAAGGACTTTCTGCACAAGGACCGtGgTGATGTCAACGTtGAGATTGTCTTT
CTTCACAAGTAAGAGCCCCCTGCTGCCACCAGACCCTGCCACC
D434G
ACTCTGGAGAGTGTCTCTAACTTCCTGAAGGACTTTCTGCACAAGGACCGtGgTGATGTCAACGTtGAGATTGTCTTTCTTCACAAGTAAGAGCCCCCTGCTGCCAC
TGAGACCTCTCACAGAGATTGAAGGACTTCCTGAAAGACGTGTTCCTGGCaCcACTACAGTTGCAaCTCTAACAGAAAGAAGTGTTCATTCTCGGGGGACGACGGTG
T L E S V S N F L K D F L H K D R G D V N V E I V F L H Exon 10
TTCCTGAAGGACTTTCTGCACAAGGACCGTGGTGATGTCAACGTTGAGATTGTCTTT
A
C
WT
TTCCTGAAGGACTTTCTGCACAAGGACCGGGATGATGTCAACGTGGAGATTGTCTTTCTTCACAAGTAAGAGCCCCCTGCTGCCACCAGACCCTGCCACCTAAACTC
AAGGACTTCCTGAAAGACGTGTTCCTGGCCCTACTACAGTTGCACCTCTAACAGAAAGAAGTGTTCATTCTCGGGGGACGACGGTGGTCTGGGACGGTGGATTTGAG
F L K D F L H K D R D D V N V E I V F L H Exon 10
Donor sequence: CTCAGAGAGAAGCATGAGTTTAGGTGGCAGGGTCTGGTGGCAGCAGGGGGCTCTTACTTcTGcAGAAAGACgAT
CTCgACGTTGACATCATCCCGGTCCTTGTGCAGAAAGTCCTTCAGG
H444Q
TTCCTGAAGGACTTTCTGCACAAGGACCGGGATGATGTCAACGTcGAGATcGTCTTTCTgCAgAAGTAAGAGCCCCCTGCTGCCACCAGACCCTGCCACCTAAACTC
AAGGACTTCCTGAAAGACGTGTTCCTGGCCCTACTACAGTTGCAgCTCTAgCAGAAAGAcGTcTTCATTCTCGGGGGACGACGGTGGTCTGGGACGGTGGATTTGAG
F L K D F L H K D R D D V N V E I V F L Q Exon 10
GACCGGGATGATGTCAACGTCGAGATCGTCTTTCTTCACAAGTAAGAGCCCCCTGC
G

Fig S 3

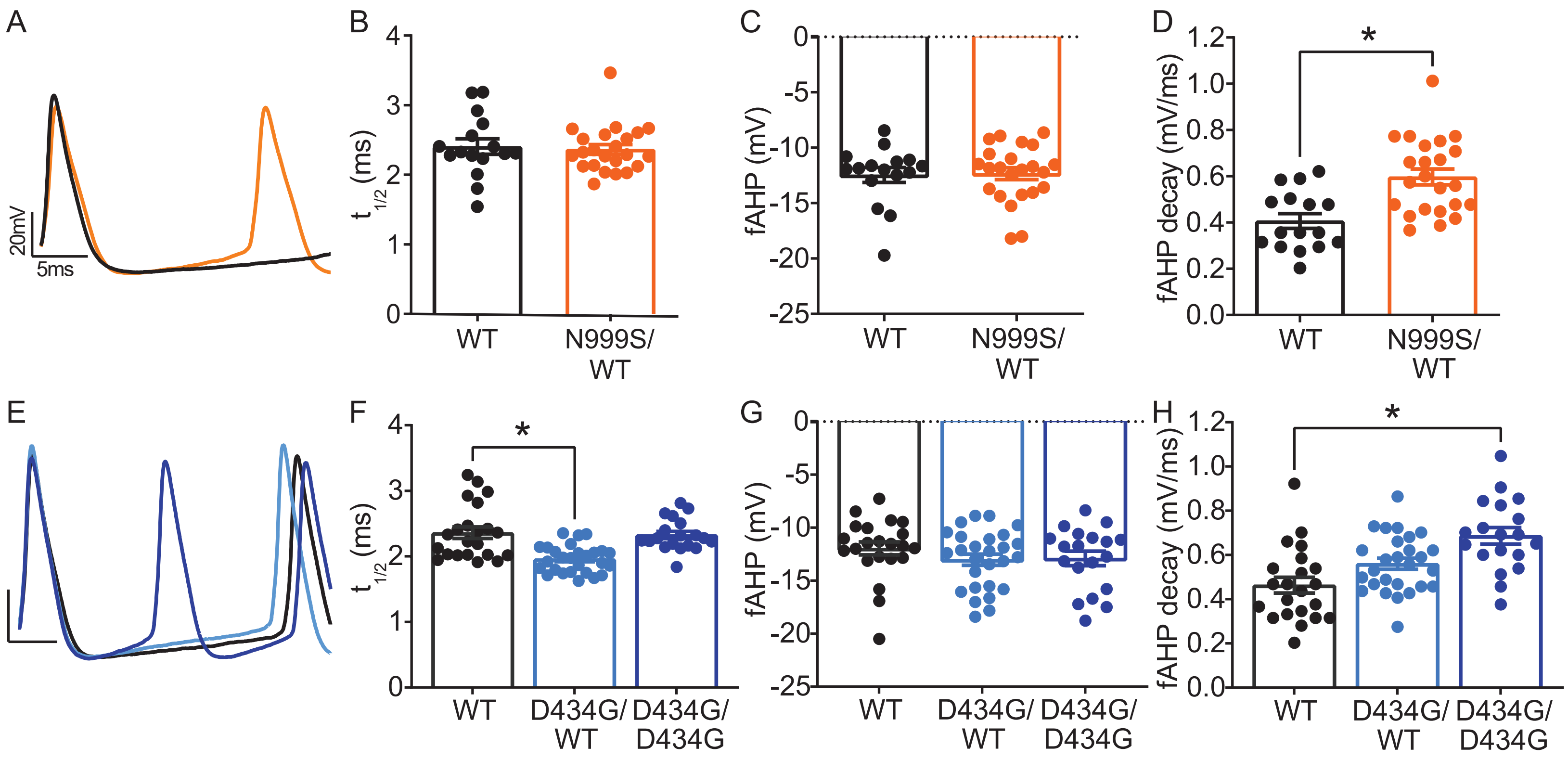

A
20mV
5ms
B
t1/2 (ms)
WT
N999S/ WT
C
fAHP (mV)
WT
N999S/ WT
D
fAHP decay (mV/ms)
*
WT
N999S/ WT
E
F
t1/2 (ms)
*
WT
D434G/ WT
D434G/ D434G
G
fAHP (mV)
WT
D434G/ WT
D434G/ D434G
H
fAHP decay (mV/ms)
*
WT
D434G/ WT
D434G/ D434G

Fig S4

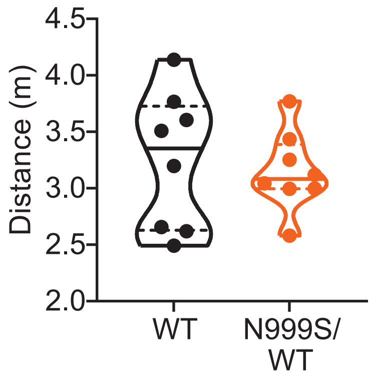

Fig S5

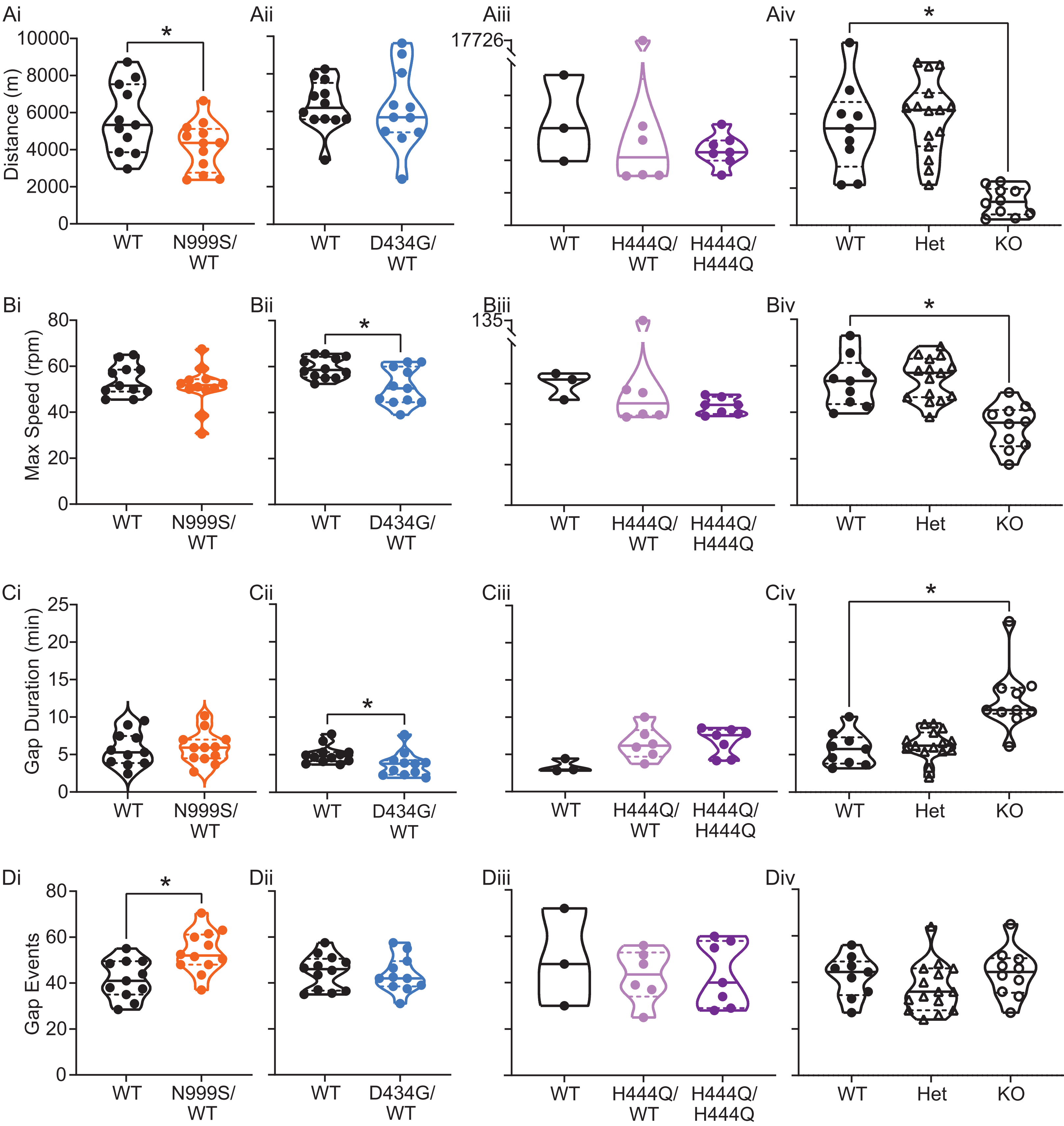

Ai
Aii
Aiii
Aiv
Distance (m)
0
2000
4000
6000
8000
10000
17726
*
WT
N999S/ WT
D434G/ WT
H444Q/ WT
H444Q/ H444Q
Het
KO
Bi
Bii
Biii
Biv
Max Speed (rpm)
0
20
40
60
80
135
Ci
Cii
Ciii
Civ
Gap Duration (min)
0
5
10
15
20
25
Di
Dii
Diii
Div
Gap Events

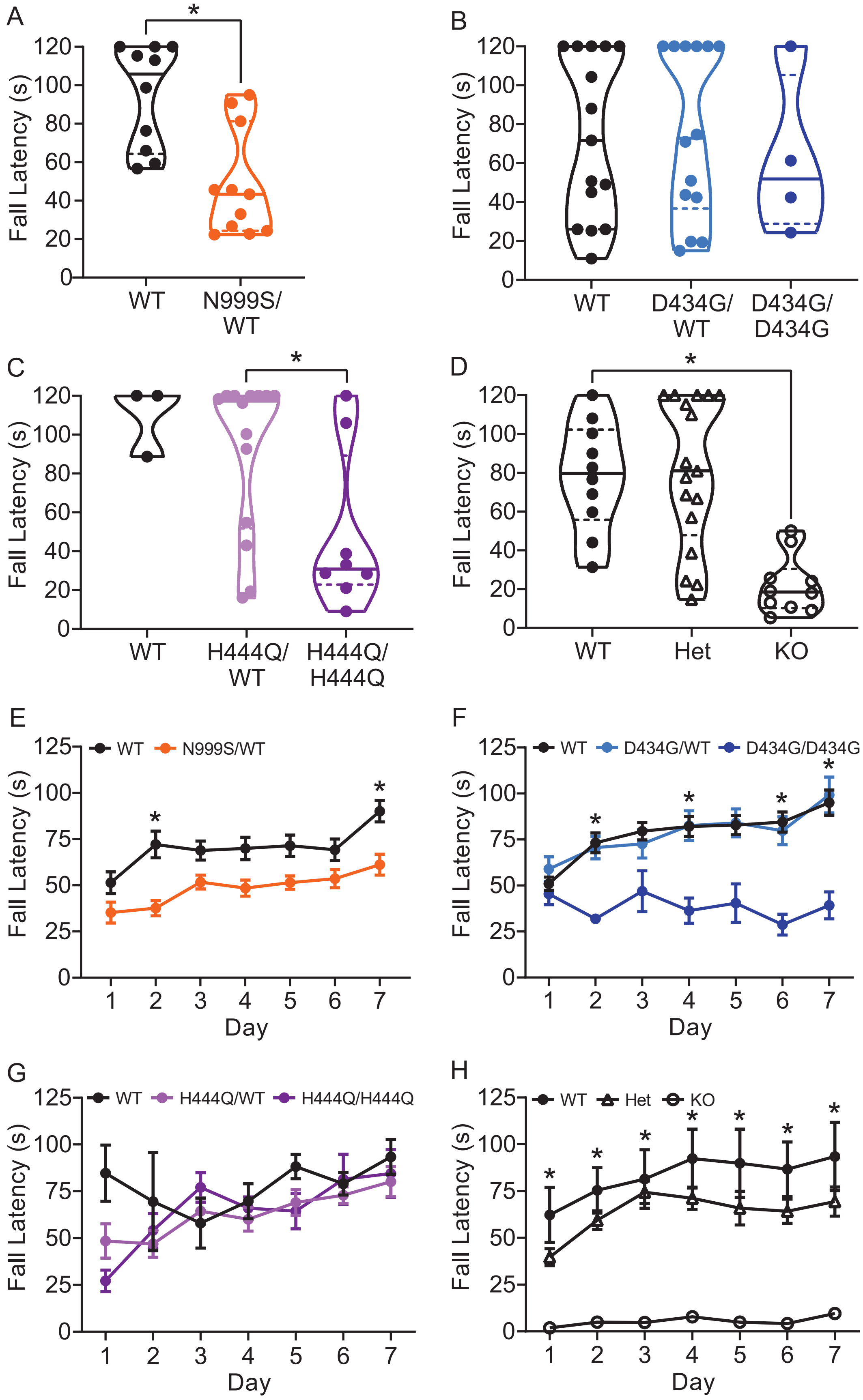
A
Fall Latency (s)
WT
N999S/WT
B
WT
D434G/WT
D434G/D434G
C
H444Q/WT
H444Q/H444Q
D
Het
KO
E
Day
WT
N999S/WT
F
D434G/WT
D434G/D434G
G
H444Q/WT
H444Q/H444Q
H
WT
Het
KO
